# Developmental Dynamics of RNA Translation in the Human Brain

**DOI:** 10.1101/2021.10.22.465170

**Authors:** Erin E. Duffy, Benjamin Finander, GiHun Choi, Ava C. Carter, Iva Pritisanac, Aqsa Alam, Victor Luria, Amir Karger, William Phu, Maxwell A. Sherman, Elena G. Assad, Alexandra Khitun, Elizabeth E. Crouch, Sanika Ganesh, Bonnie Berger, Nenad Sestan, Anne O’Donnell-Luria, Eric Huang, Eric C. Griffith, Julie D. Forman-Kay, Alan M. Moses, Brian T. Kalish, Michael E. Greenberg

## Abstract

The precise regulation of gene expression is fundamental to neurodevelopment, plasticity, and cognitive function. While several studies have deeply profiled mRNA dynamics in the developing human brain, there is a fundamental gap in our understanding of accompanying translational regulation. We perform ribosome profiling from more than 70 human prenatal and adult cortex samples across ontogeny and into adulthood, mapping translation events at nucleotide resolution. In addition to characterizing the translational regulation of annotated open reading frames (ORFs), we identify thousands of previously unknown translation events, including small open reading frames (sORFs) that give rise to human- and/or brain-specific microproteins, many of which we independently verify using size-selected proteomics. Ribosome profiling in stem cell-derived human neuronal cultures further corroborates these findings and shows that several neuronal activity-induced long non-coding RNAs (lncRNAs), including *LINC00473*, a primate-specific lncRNA implicated in depression, encode previously undescribed microproteins. Physicochemical analysis of these brain microproteinss identifies a large class harboring arginine-glycine-glycine (RGG) repeats as strong candidates for regulating RNA metabolism. Moreover, we find that, collectively, these previously unknown human brain sORFs are enriched for variants associated with schizophrenia. In addition to significantly expanding the translational landscape of the developing brain, this atlas will serve as a rich resource for the annotation and functional interrogation of thousands of previously unknown brain-specific protein products.

## MAIN

The human brain leverages extraordinary protein diversity to execute developmental programs, organize neural circuits, and perform complex cognitive tasks ^1^. Proteomic diversity is generated through a series of transcriptional, post-transcriptional, and translational mechanisms that ultimately contribute to a rich and complex ‘translatome’. While many studies have focused on genomic and transcriptomic regulation in the developing human brain, much less is known regarding the complexity of translational regulation in this context, underscoring the need to study this key regulatory node in human brain development.

Deep sequencing of ribosome-protected mRNA fragments (ribosome profiling) provides a means to map genome-wide translation at nucleotide resolution ^2^. From these data, the movement of ribosomes across codons can be determined and then used to identify protein-coding open reading frames (ORFs). Ribosome profiling in various systems has revealed that the fraction of the transcriptome subject to translation is far greater than previously recognized, with a single transcript often encoding many distinct protein products. Thus, RNA-seq analysis fails to give a complete picture of the landscape of proteins produced in the brain. Indeed, studies in yeast ^3^, as well as cardiac ^4^ and tumor tissues ^5^, have revealed the widespread active translation of small ORFs (sORFs) encoding microproteins ≤100 amino acids. From the relatively few microproteins to be functionally investigated so far, researchers have identified important regulators of mitochondrial metabolism, translational regulation, and cell differentiation ^6–8^. To date, however, the nature and roles of microprotein species in the developing human brain remain almost entirely uncharacterized.

Here we describe a comprehensive translational atlas of the human prenatal and adult cortex, involving more than 70 distinct tissue samples. In addition to cataloguing annotated gene programs subject to dynamic regulation at the translational level, we identify a vast array of novel sORFs and other non-canonical translation events, including many arising from previously annotated non-coding transcripts. We subsequently employ size-selected proteomics to independently verify a subset of these products at the protein level. Similar findings were also obtained upon ribosome profiling of stem cell-derived human neuronal cultures, where we identify several novel microproteins translated from neuronal activity-responsive RNAs previously annotated as non-coding, including *LINC00473*, a primate-specific lncRNA previously implicated in depression.

Notably, nearly one in five of our newly identified sORFs derive from brain-enriched or brain-specific transcripts, suggesting that their functions may be unique to the brain. Among these, we identify hundreds of microproteins that are functionally related to the RGG domain of intrinsically disordered RNA-binding proteins, suggesting that these protein products may modulate RNA metabolism and/or function. We find that the majority of sORFs in the brain are newly evolved in humans, where a subset of sORFs arose via transposable element insertion at start codons. While their recent evolution might be thought to suggest that the microproteins translated from these sORFs are non-functional, we identify >100 human-specific microproteins that play a key role in cell viability.^9^ Moreover, we find that human brain sORFs are significantly enriched for schizophrenia disease heritability, suggesting that microproteins encoded by these sORFs may play a significant role in disease etiology. Our study thus significantly expands the known translational landscape of the developing brain and provides a rich resource for the study of novel human brain sORFs, which is accessible via our accompanying web-based searchable database (http://greenberg.hms.harvard.edu/project/human-brain-orf-database/).

## RESULTS

### Translational Landscape of the Human Prenatal and Adult Brain

To characterize the human brain translational landscape at single-nucleotide resolution, we performed simultaneous RNA sequencing (RNA-seq) and ribosome profiling (Ribo-seq) from human adult dorsolateral prefrontal cortex and prenatal cortex across a range of ages (Fig. 1a). RNA-seq provides a quantitative measure of the mRNA species expressed in the brain, whereas Ribo-seq allows for a quantitative appraisal of active mRNA translation. The gestational age of prenatal cortex samples (30 total) ranged from 12 to 23 weeks, while adult brain donors (43 total) ranged in age from 18 to 82 years, with an average post-mortem interval of 9.9 hours (Fig. 1b). Importantly, across samples, Ribo-seq data exhibited the three-nucleotide periodicity characteristic of actively translating ribosomes, a key metric for confident ORF identification (Fig. 1c). Moreover, Ribo-seq reads exhibited expected fragment size distributions (Fig. S1b) and mapped primarily to annotated gene coding regions (Fig. S1c), further supporting the idea that this method robustly captures RNA protected by actively translating ribosomes. Full demographic and Ribo-seq quality metrics are available in Table 1 and Figure S1.

**Figure 1:**
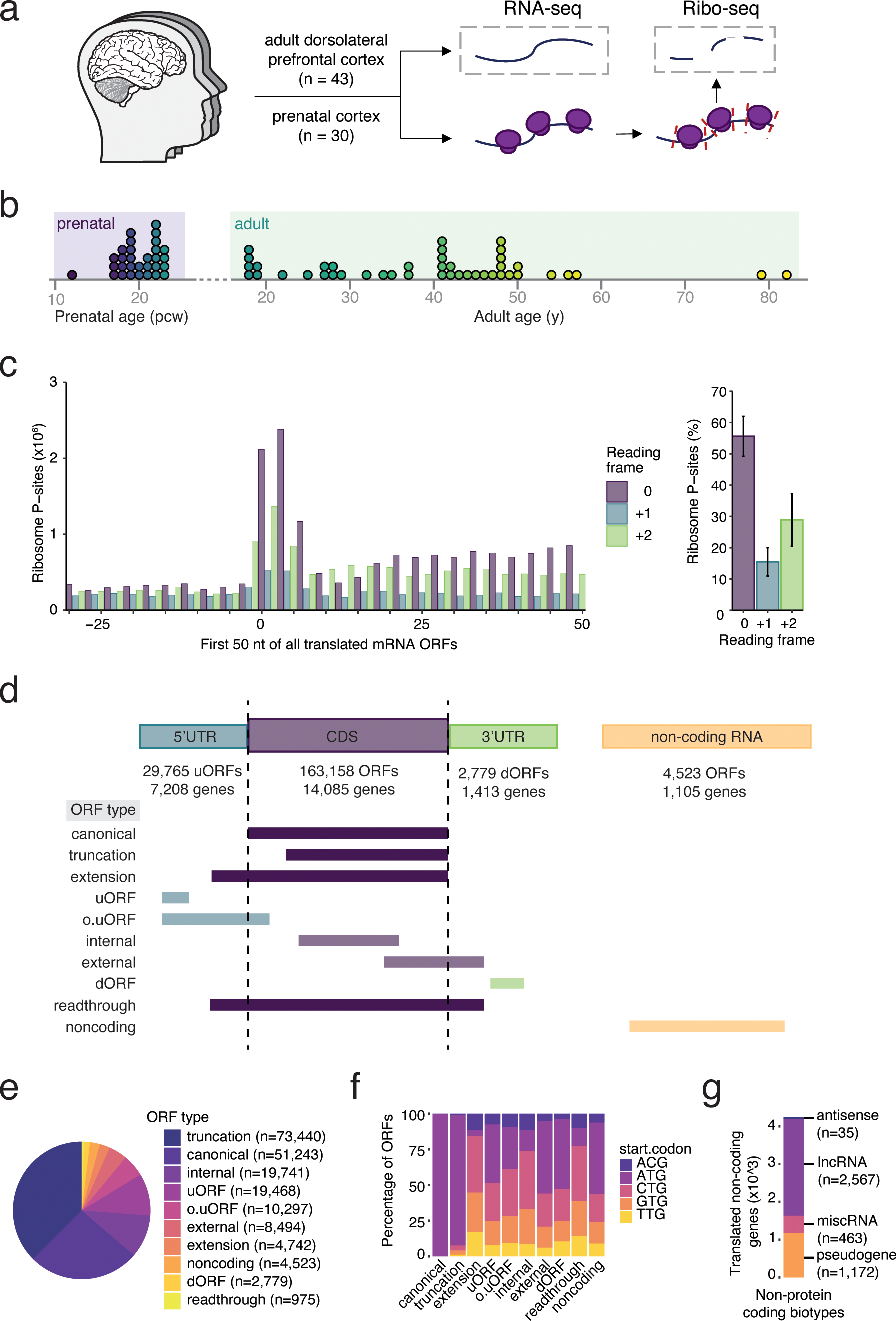
Ribosome profiling captures active translation in the human adult and prenatal brain. (a) Overview of experimental design. (b) Histogram depiction of patient samples included in this study. (c) Bar plot displaying P-sites derived from offset-corrected Ribo-seq reads in the first 100nt of annotated ORFs (left) and the percentage of footprints in each reading frame (right). (d) Schematic overview of ORF types detected by RibORF. (e) Number of ORFs of each type identified in human adult and/or prenatal brain. (f) Stacked bar plot of start codon usage by ORF type. (g) Stacked bar plot of numbers and percentages of translated non-coding RNAs separated by transcript biotype.

High-confidence bona fide ORFs were identified based on the characteristic triplet reading frame periodicity of ribosome footprints using RibORF ^10^, with sequences present in two or more samples, exhibiting clear start and stop codons, and displaying Ribo-seq reads across the entire putative ORF region. After combining data across samples and filtering for ORF quality, we identified a total of 195,702 distinct actively translated ORFs in the human brain, mapping to 14,234 distinct genes (Fig. 1d & e, Fig. S1d). In support of the quality of the resulting annotations, the relative proportions of each ORF type in our dataset, as well as general features such as start codon usage, were broadly consistent with previous findings in cell lines ^11, 12^ and other tissues ^13^ (Fig. 1e, f). ORFs translated from non-coding RNAs were most commonly identified within previously annotated lncRNAs or pseudogenes (Fig. 1g), including the recently characterized lncRNA-encoded microproteins NoBody ^8^, MOXI ^6^, and Cyren ^14^. However, other highly expressed lncRNAs that are not known to be translated such as *XIST*, *HOTAIR*, and *NEAT1* showed no evidence of active translation in the brain, further corroborating the specificity of the identified lncRNA-associated ORFs. Taken together, these data map the translational landscape of the human cortex across development at an unprecedented level of resolution.

### Transcriptional and Translational Regulation of Human Brain Development

While transcriptional changes during neurodevelopment have been extensively profiled ^15, 16^, the contribution of translational regulation during neurodevelopment is still poorly understood. Adopting previous methods that use the number of ribosomes per RNA molecule (ribosome density) as a measure of translational efficiency, we first investigated the extent to which brain ORFs exhibit developmental shifts in translational efficiency ^17^, focusing on canonical ORFs that encode proteins of known function. Indeed, comparison of our paired transcriptome and translatome datasets revealed several distinct modes of developmental regulation (Fig. 2a & b, S2). In this regard, we found, for example, that developmental decreases in ribosomal gene RNA levels were effectively buffered by corresponding increases in translational efficiency (Fig. 2c, S2a & b). This is consistent with previous reports that ribosomal genes exhibit increased DNA methylation, an epigenetic silencing mark, during postnatal development ^18^, and suggests that a compensatory increase in translation of these genes may be required to maintain proteostasis in the developing brain. In contrast, ORFs encoding both major histocompatibility complex (MHC) components and proteins involved in complement activation exhibited synergistic increases in both mRNA levels and translational efficiency between the prenatal and adult brain (Fig. 2c, S2a & b). Given the respective roles of these factors in developmental synapse formation and elimination ^19, 20^, these findings implicate active translational regulation in the control of developmental synaptic pruning and circuit assembly. Further investigation of these pervasive forms of translational regulation promises new insights into the gene expression mechanisms that control various aspects of human neurodevelopment.

**Figure 2:**
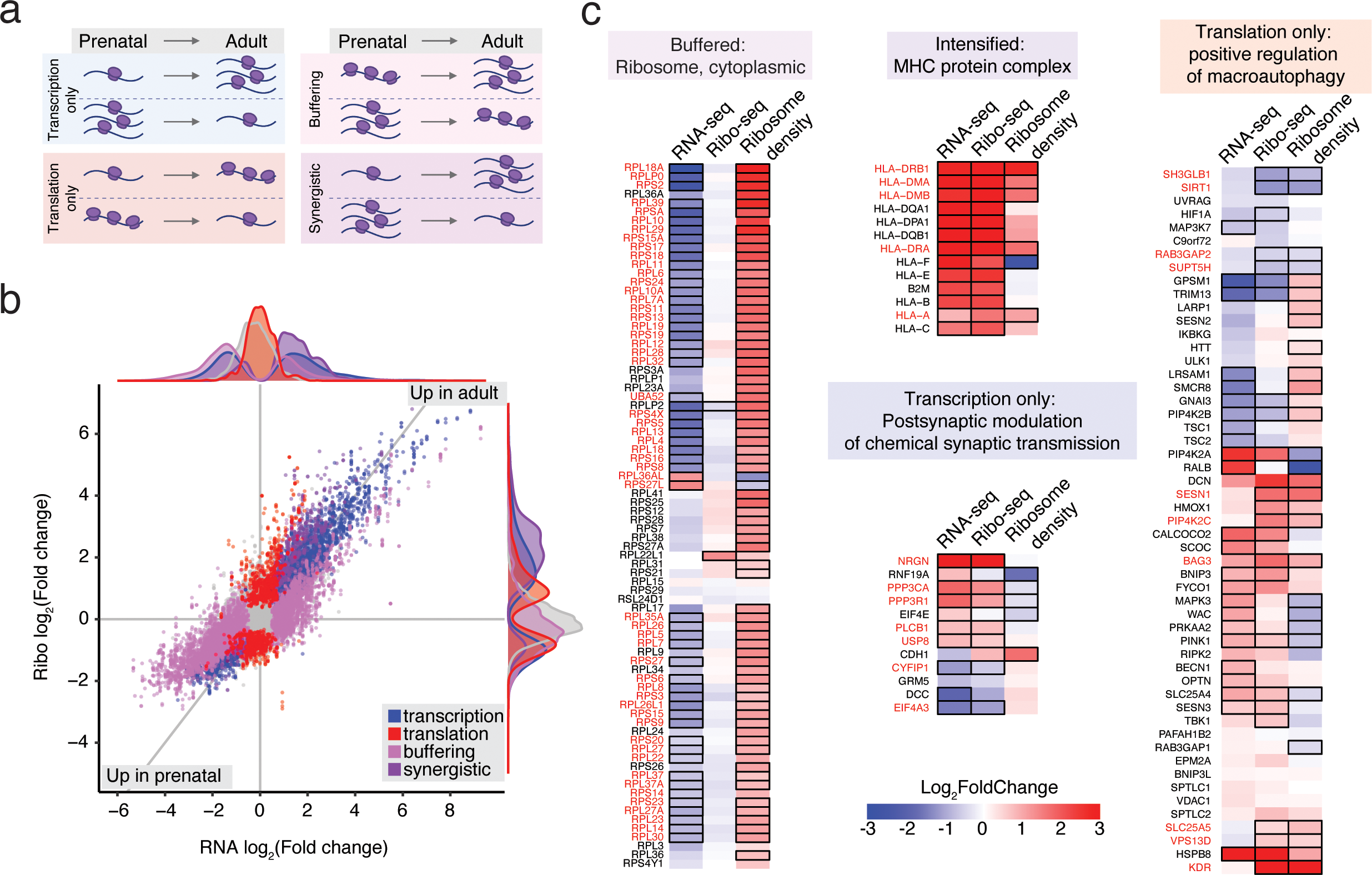
Transcriptional and translational regulation across human brain development. (a) Classification of genes based on RNA-seq, Ribo-seq, and ribosome density measurements. (b) Scatterplot of fold-changes between adult and prenatal brain for all canonical ORFs in Ribo-seq data and the corresponding gene in RNA-seq data. Positive values indicate enrichment in the adult brain, whereas negative values indicate enrichment in the prenatal brain. Transcriptionally forwarded genes (blue; change in transcription with no change in ribosome density), translationally exclusive genes (red; change in ribosome density with no change in transcription), buffering genes (light purple; change in ribosome density that counterbalances the change in mRNA transcription), and synergistic genes (dark purple; change in ribosome density that amplifies the change in mRNA) are highlighted. (c) Heatmap of genes associated with the top GO term in each regulatory category identified in A. Black outlines indicate p < 0.05, gene names in red indicate inclusion in a given regulatory category.

### sORFs and Non-canonical Translation in the Human Brain

Previous studies in other systems have shown that translational regulation is more widespread across the transcriptome than previously appreciated, often involving regions of the genome previously annotated as non-coding (e.g. pseudogenes, lncRNAs, antisense RNAs, and 5’ and 3’ untranslated regions of canonical protein-coding genes). To interrogate our datasets for novel human brain microproteins, we focused on Ribo-seq-identified ORFs ≤300 nucleotides (nt) (100 AAs) in length that were either out of frame or did not overlap with longer ORFs. This analysis identified 45,109 actively translated sORFs originating from 9,219 genes in the prenatal or adult brain (Fig. 3a, S3a & b). While many of these sORFs were translated from alternative regions of canonical protein-coding transcripts, 3,132 were derived from annotated non-coding transcripts, including reported lncRNAs, pseudogenes, and antisense transcripts (Fig. 3b). Importantly, while the ribosome density of sORFs was on average ∼10-fold lower than the translation of canonical ORFs (Fig. S3a), this was also true of a number of previously reported microproteins with well-characterized functions, including RPL41 ^21^, SLN ^22^, and NBDY ^8^, suggesting that newly described sORFs with relatively low translation compared to canonical ORFs are likely to be functional. Like canonical ORFs, many of these sORFs were developmentally regulated via coordinated changes in RNA abundance and/or translational efficiency, which likely enables the fine-tuning of sORF protein levels as the brain matures (Fig. 3c).

**Figure 3:**
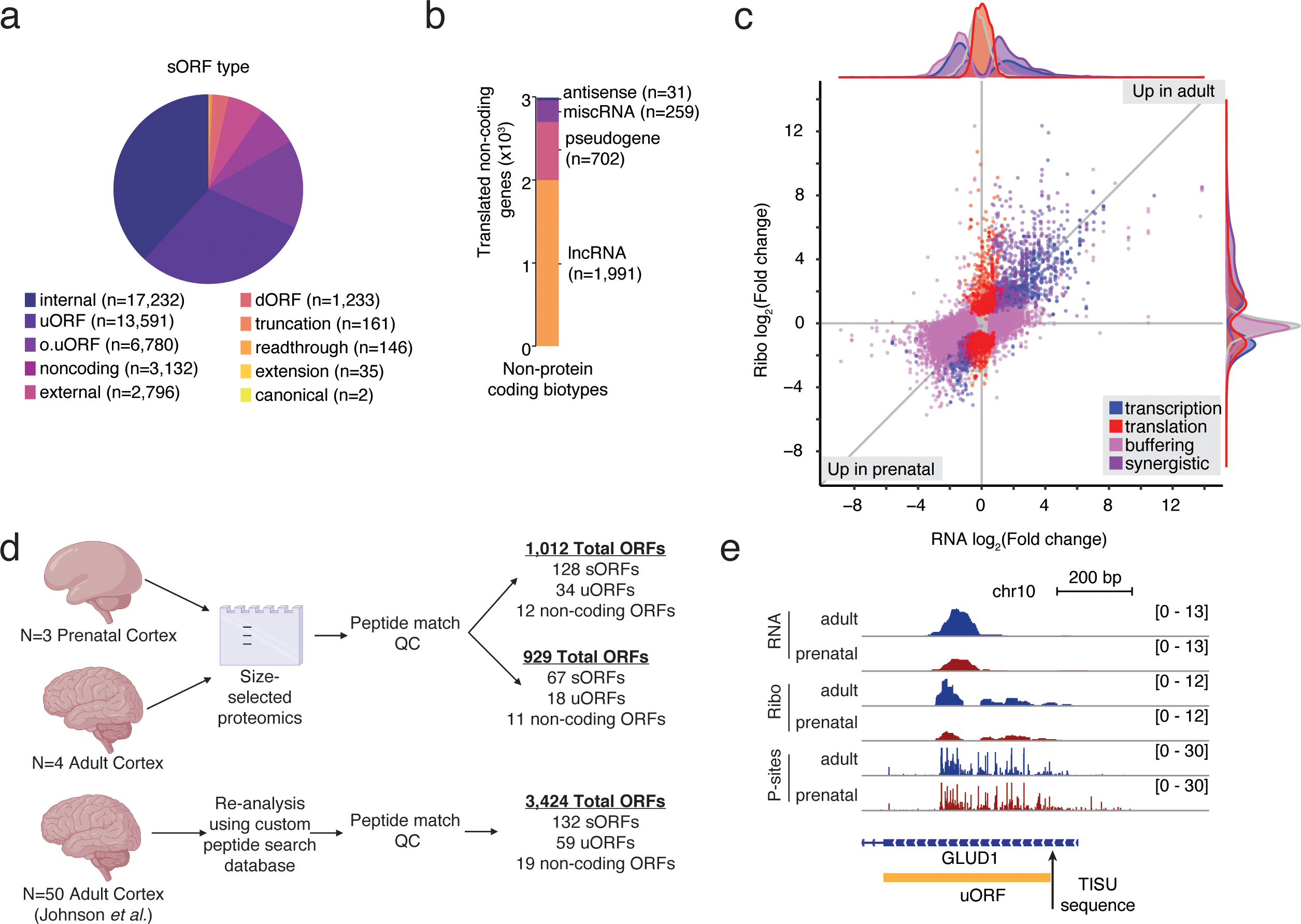
Microprotein expression and validation across brain development. (a) Number of sORFs of each type identified in human adult and/or prenatal brain. (b) Stacked bar plot of numbers and percentages of translated non-coding RNAs containing at least one sORF, separated by transcript biotype. (c) Scatterplot of fold-changes between adult and prenatal brain for all sORFs in Ribo-seq data and the corresponding gene in RNA-seq data. Positive values indicate enrichment in the adult brain, whereas negative values indicate enrichment in the prenatal brain. Translationally forwarded genes (blue), exclusive genes (red), buffering genes (light purple), and synergistic genes (dark purple) are highlighted. (d) Number and type of ORFs identified by size-selection proteomics in the adult and prenatal brain, or by Johnson *et al.* (b) Number of ORFs binned by protein length identified by size-selection proteomics in the adult and prenatal brain, or by Johnson *et al.* (e) Genome locus of *GLUD1.* Tracks represent merged and depth-normalized reads across all adult vs. prenatal samples for RNA-seq, Ribo-seq, as well as P-site positions. The sORF identified by RibORF is shown in gold.

While recognizing the difficulties associated with proteomic microprotein detection, we sought to independently corroborate our Ribo-seq findings at the protein level. Towards this end, we performed size-selected mass spectrometry-based proteomics for enhanced detection of protein species less than 20 kDa ^23^. To facilitate the identification of proteins not annotated by Uniprot, this analysis incorporated a proteogenomic approach, whereby all peptides detected by mass spectrometry were matched to a custom database constructed from our Ribo-seq data (See Extended Methods). To further increase our ability to detect rare microproteins, we also reanalyzed published mass spectrometry data from 50 human adult brain tissue samples to search for signatures of sORF-derived microproteins ^24^.

Collectively, these analyses identified peptides corresponding to 4,224 unique ORFs (Fig. 3d, S3c-e), including 239 sORFs, 13 of which derived from previously annotated non-coding transcripts. Given the challenges associated with proteomic microprotein discovery, these results constitute a powerful independent validation of our Ribo-seq findings. Moreover, these biochemically verified microproteins represent strong candidates for further investigation. To highlight one such example, this analysis confirmed the presence of a novel microprotein encoded by a upstream ORF (uORF) in *GLUD1* (*Glutamate dehydrogenase 1*), a gene critically involved in glutamate metabolism (Fig. 3e) ^25, 26^. Notably, this *GLUD1* uORF contains a Translation Initiator of Short 5′ UTR (TISU) motif, which is known to enable uninterrupted translation under conditions of energy stress, a condition that leads to translational inhibition of most other mRNAs ^27^. This suggests that this microprotein might contribute to neuronal responses to acute metabolic demands. A full list of proteomically validated sORF species is available in Table 2. These proteogenomic datasets validate the presence of a large number of non-canonical ORF-derived microproteins that are expressed in the brain at levels similar to microproteins that have been identified in other tissues and found to have critical biological functions. These microproteins represent a significant expansion of the known brain translatome with potentially immense functional relevance for human development and disease.

### Regulated sORF Translation in Human Neurons

To complement these tissue-based studies, we also characterized the translational landscape in human embryonic stem cell (hESC)-derived neuronal cultures. To this end, we employed an engineered hESC line harboring an integrated doxycycline-inducible NGN2 construct, treating with doxycycline after plating to induce NGN2 expression. Adapting a previously described protocol ^28^, this approach was combined with SMAD and WNT inhibition to induce patterning toward a forebrain phenotype (see Extended Methods; Fig. 3a). The resulting cultures (hereafter NGN2 neurons) demonstrate transcriptional signatures largely consistent with well-differentiated glutamatergic neurons (Fig. S3a).

For these studies, we also exploited the ability to induce acute, synchronous membrane depolarization in this system to examine neuronal activity-responsive translational changes. Thus, day 28 cultures from three independent differentiation cohorts were harvested for combined RNA-seq and ribosome profiling either prior to or 6 h following depolarization with 55 mM potassium chloride (KCl). The resulting datasets passed key quality control metrics, with clear three-nucleotide periodicity observed in the Ribo-seq data (Fig S4b) and high data correlation between separate differentiation cohorts (Fig. S4c & d). Moreover, robust induction of known activity-responsive loci was observed in all depolarized samples (Fig. S4e & f).

Collectively, this analysis identified a total of 106,792 actively translated ORFs in NGN2 neurons (Fig 4b & c, S4g), >50% of which (59,404/117,992) were also observed in the human brain tissue samples. As expected, however, principal component analysis (PCA) plots showed that NGN2 samples cluster more with the fetal than adult translatome (Fig 4d).

**Figure 4:**
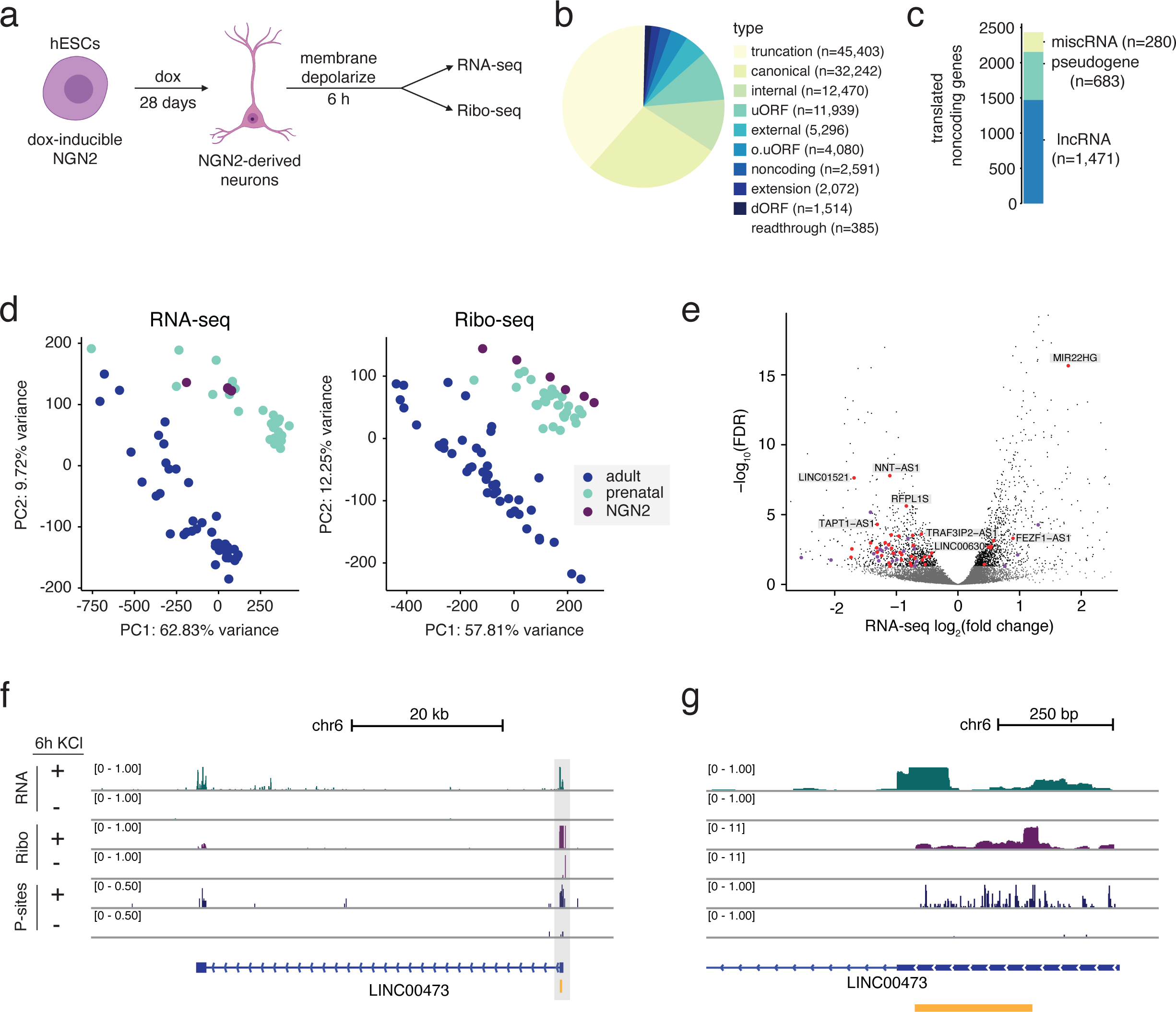
Activity-dependent translation in hESC-derived neurons. (a) Schematic of Ribo-seq and RNA-seq from NGN2-derived hESCs following 6h membrane depolarization. (b) Breakdown of translated ORFs of each type identified in NGN2-derived neurons. (c) Stacked bar plot of numbers and percentages of translated non-coding RNAs separated by transcript biotype. (d) PCA analysis based on RNA-seq and Ribo-seq reads mapping to annotated genes in primary adult and prenatal brain tissue and NGN2 neurons. (e) Volcano plot of -log_10_(p_adj_) versus log_2_(fold-change) in RNA-seq expression between membrane-depolarized and unstimulated NGN2 neurons. Black indicates p_adj_ < 0.05, purple indicates activity-dependent non-coding RNAs with no evidence of translation in human brain or NGN2 neurons, red indicates activity-dependent non-coding RNAs with evidence of translation in human brain and/or NGN2 neurons. (c) Genomic locus of *LINC00473* in NGN2 neurons. Tracks represent merged and depth-normalized reads across 3 biological replicates of membrane-depolarized (6 h KCl) and unstimulated neurons for RNA-seq, Ribo-seq, as well as P-site positions. sORFs identified by RibORF are shown in gold.

Given this broad translational overlap, we focused our attention on novel sORFs translated from previously annotated non-coding RNAs (ncRNAs). In this regard, we observed active translation of novel sORFs in 39 of 78 ncRNAs detected in NGN2 cultures (Fig. 4e, S4h-j), many of which (25) were also detected by Ribo-seq in our human brain datasets. Moreover, a number of the resulting protein products could be verified biochemically using size-selected proteomics (Fig. S4k, Table 2). Among our findings, we uncovered an actively translated sORF within *LINC00473* (Fig 4f & g), which we previously identified as a primate-specific and activity-dependent lncRNA ^29^ and has been implicated as a sex-specific driver of stress resilience when expressed ectopically in the mouse prefrontal cortex ^30^. Thus, neuronal activity-dependent expression of annotated non-coding transcripts is frequently associated with the unappreciated translation of novel microproteins, which may modulate key neuronal responses to activity.

### Evolutionary Conservation of sORFs

To begin to investigate possible human brain sORF function more broadly, we analyzed the evolutionary origins of brain sORFs using genomic phylostratigraphy – an approach that dates the origin of individual genes by examining the presence or absence of homologs across species (see Extended Methods) ^31^. Determination of the minimal evolutionary age for human brain sORFs revealed that, in contrast to most annotated protein-coding genes, a majority of sORFs are human-specific (12% and 65%, respectively Fig. 5a, S5a). This analysis further revealed that more recently evolved sORFs are shorter, contain fewer splice junctions, and exhibit lower ribosome density compared to their more evolutionarily ancient counterparts (Fig. 5b-d). Microproteins encoded by more evolutionarily ancient sORFs are also more likely to be detectable by proteomics, perhaps indicative or more overall higher levels of expression (Fig. S5b). These features are consistent with the classic view that more evolutionarily conserved regions of the genome are more likely to be functional sequences, nominating highly conserved sORFs as promising candidates for functional studies. However, the rapid evolution of human-specific sORFs also suggests that these sequences may represent evolutionary experiments - regions that gain translation capacity in any given lineage that may not always be conserved during further evolution. To test whether a subset of newly-evolved sORFs may be functional, we overlapped sORFs that show evidence of translation in the human brain with a recently published dataset of CRISPR-Cas9 perturbation of sORFs in K562 or iPSC human cells, selecting for those sORFs that show a significant growth phenotype upon knockout (Mann-Whitney U p-value <0.05).^9^ Of the 139 sORFs that satisfy these criteria, a striking 113 are human-specific, which lends strong support to the hypothesis that newly evolved microproteins can exhibit important functions in humans. Moreover, it is also notable that, relative to all sORFs, those derived from brain-enriched transcripts are significantly more likely to be specific to humans (K-S test, P<2.2e-16, Fig. S5c), which is consistent with the idea that these protein products might contribute to human-specific aspects of brain development.

**Figure 5:**
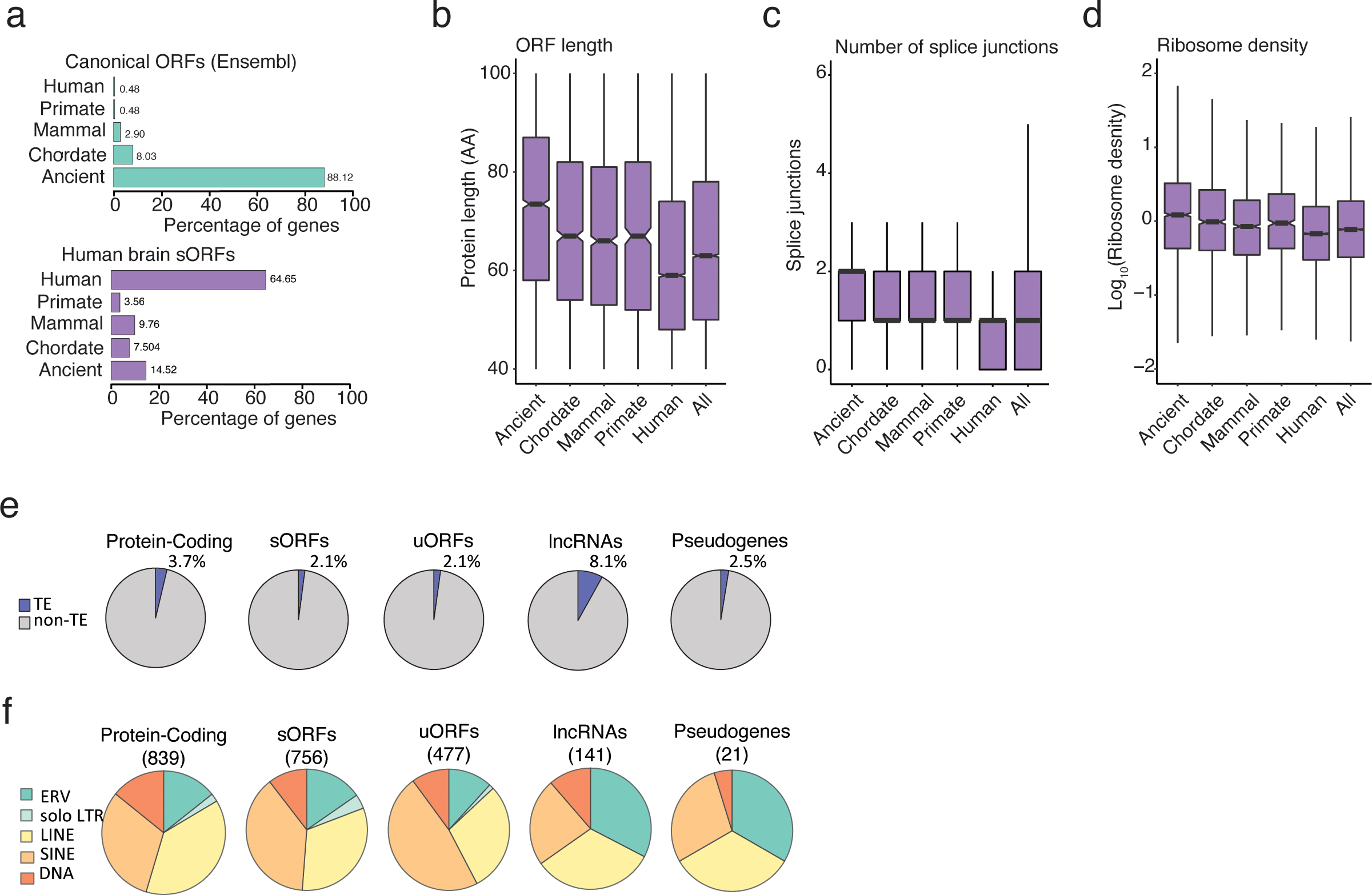
Evolutionary origins of human brain sORFs. (a) Number and percentage of canonical ORFs (top, all ORFs in human Ensembl database, ≥40 AA) and sORFs (bottom, ≥40 AA) grouped by evolutionary age. (b) Box and whisker plots of microprotein ORF length grouped by evolutionary age. (c) Box and whisker plots of the number of splice junctions per microprotein ORF (40-100 AA) grouped by evolutionary age. (d) Box and whisker plots of microprotein ORF ribosome density grouped by evolutionary age. (e) Pie chart of the percentage of ORFs with a TE insertion at the start codon, grouped by ORF type or non-coding RNA biotype. (f) Pie chart of the distribution of TE types, grouped by ORF type or non-coding RNA biotype. Numbers indicate the number of ORFs in each category.

Recently, Playfoot and colleagues provided evidence for transposable element (TE) involvement in new ORF formation ^32^. We directly explored this as a possible mechanism of sORF generation in the brain, finding that lncRNA-associated sORFs have an increased overlap with TE insertions compared to protein-coding ORFs (8% vs. 4%, respectively; Fig. 5e, S5e-f). This TE enrichment within lncRNAs has been previously noted and suggested to contribute new non-coding sequences for RNA-mediated lncRNA function ^33–35^. Our findings, however, provide evidence that TEs might also play an important role in the generation of new protein-coding ORFs within these annotated non-coding regions. Notably, different classes of TEs were also found to be associated with distinct ORF types (Fig 5f); however, the function significance of this observation awaits further investigation.

### Upstream ORF (uORF) Regulation of Canonical Protein Translation

Of the actively translated sORFs identified from the human brain, 8,446 (19%) were translated from brain-enriched or brain-specific transcripts ^36^, suggesting that in many cases their functions may be unique to the brain. To more directly investigate sORF function, we first focused on uORFs, a category of ORFs commonly thought to negatively regulate downstream translation of canonical ORFs through a variety of mechanisms, including stalled translational termination ^37, 38^. Somewhat surprisingly, but consistent with more recent findings ^4, 37–39^, we found that uORF translation was not generally anti-correlated with translation of the corresponding canonical ORF (Fig. 6a-c, S6a-c). Notwithstanding this general finding, we still identified several individual uORFs that were strongly anti-correlated with translation of their canonical downstream ORFs. One such example involved a uORF in *DLGAP1*, which encodes an important brain-enriched post-synaptic scaffolding protein (Fig. 6d-f). In this case, translation of the *DLGAP1* uORF was strongly enriched in prenatal samples via the preferential use of an alternative transcriptional start site (Fig. 3d & e, S3d), and increased uORF translation is associated with a reduction in translation of the canonical *DLGAP1* protein (Fig. 3f). Together, these data point toward a mechanism in which the use of an alternative TSS in the prenatal, but not the mature, brain leads to specific translational repression of the canonical *DLGAP1* protein. Notably, *DLGAP1* is a known autism-associated gene ^40^, raising the possibility that this developmentally-timed regulation of *DLGAP1* translation may be required for proper neurological function. Our overall findings are thus consistent with a nuanced role for brain uORFs in translational regulation, with select uORFs exerting a strong negative regulatory influence on developmentally timed protein expression.

**Figure 6:**
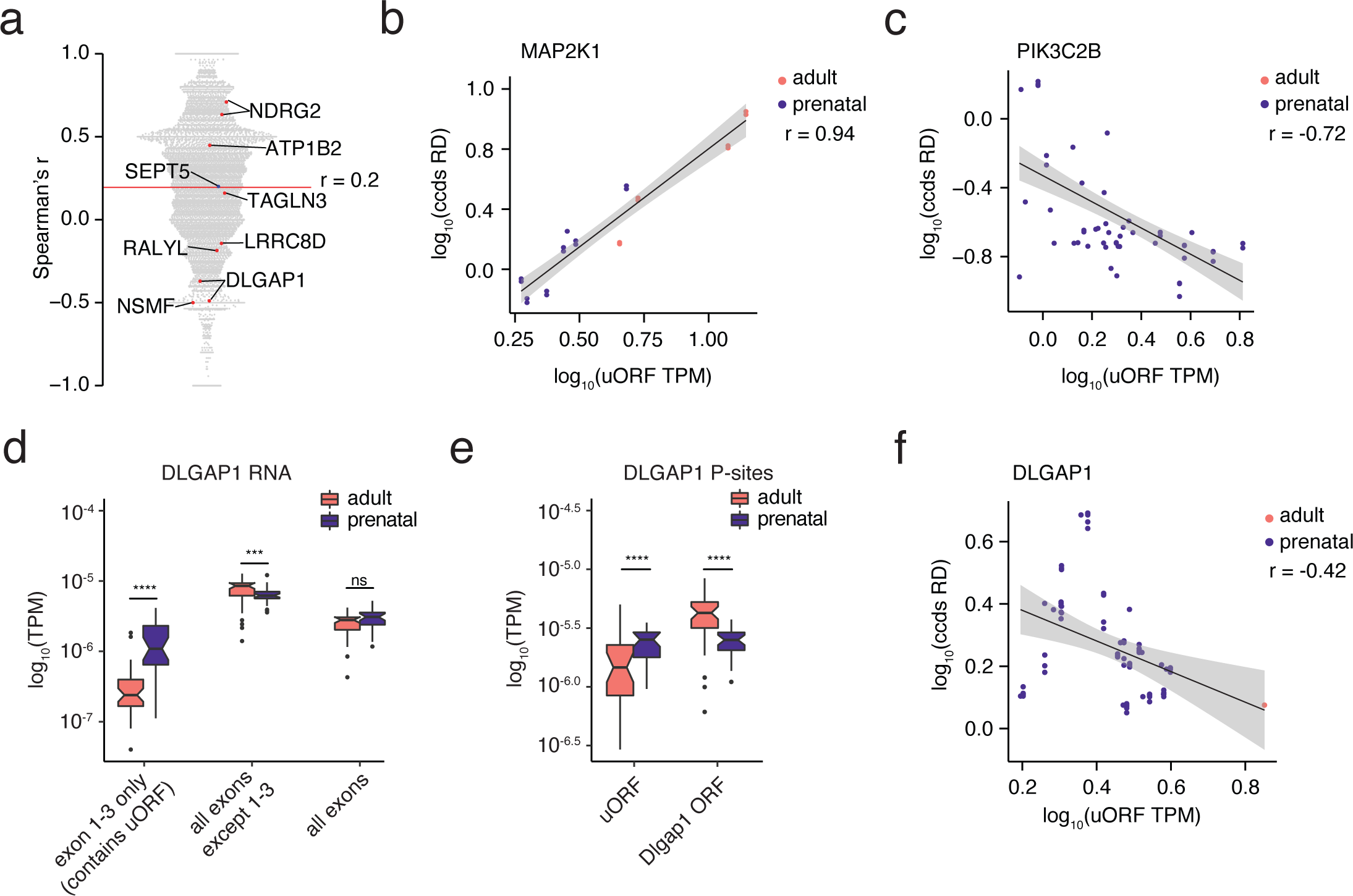
Effects of uORF expression on downstream ORF translation. (a) Beeswarm dot plot showing the Spearman’s r correlation between uORF occupancy and canonical ORF TE for individual genes across all 73 individuals. Red line represents the mean correlation across all genes. Red dots indicate developmentally regulated uORFs (described in Fig. S6a). (b & c) Scatterplot and Spearman’s r correlation between upstream ORF translation (uORF TPM) and canonical ORF ribosome density (ccds TPM) for *MAP2K1* (b) and *PIK3C2B* (c) across 73 individuals. (d) Box and whisker plot of RNA-seq reads from adult and prenatal samples over *DLGAP1* exons 1-3, all exons except 1-3, and all exons. (e) Box and whisker plot of Ribo-seq P-sites from adult and prenatal samples over *DLGAP1* uORF and ccds ORF. **** p < 0.0001, ***p < 0.001, KS test. (f) Scatterplot and Spearman’s r correlation between upstream ORF translation (uORF TPM) and canonical ORF translation (ccds TPM) for *DLGAP1* across 73 individuals.

### Sequence-based Physicochemical Analysis of Brain Microprotein Function

Beyond general evolutionary considerations, we also sought to gain further insight into brain microprotein function through primary sequence analysis. In this regard, 7,471 (17%) human brain sORFs showed significant sequence similarity (E < 10^-4^) to known proteins, with 699 (∼2%) matching a protein sequence encoded elsewhere in the genome. These previously characterized protein paralogs participate in a variety of processes, including cellular metabolism, transcription, translation, and membrane transport (Fig. S7a, Table 3), raising the possibility that their corresponding sORFs encode microproteins with related biological functions. Indeed, we found that 34% of the sORFs with significant sequence similarity to known or predicted human proteins overlapped with an annotated protein domain, strongly suggesting that many of these sORFs encode defined folded structures or even entire structural domains (Fig. S7b).

For sORFs lacking sequence similarity to known or predicted human proteins (67%), calculated FoldIndex scores, a rough predictive measure of intrinsic disorder ^41^, suggest that these protein products do not generally adopt stable three-dimensional conformations (Fig. 7a). Consistent with this idea, overall, sORFs exhibited significantly (all *P*<10^-10^, T-test) reduced sequence complexity (W-F complexity 0.68 vs. 0.76), lower aromaticity (5.4 vs. 8.3 aromatic residues per 100 AA), higher isoelectric point (8.8 vs. 8.0), and elevated arginine to lysine ratios (log ratio 1.1 vs. -0.05) relative to the known human proteome, all indicative of the absence of well-defined three-dimensional structure (Fig. 7b, S7c-h).

**Figure 7:**
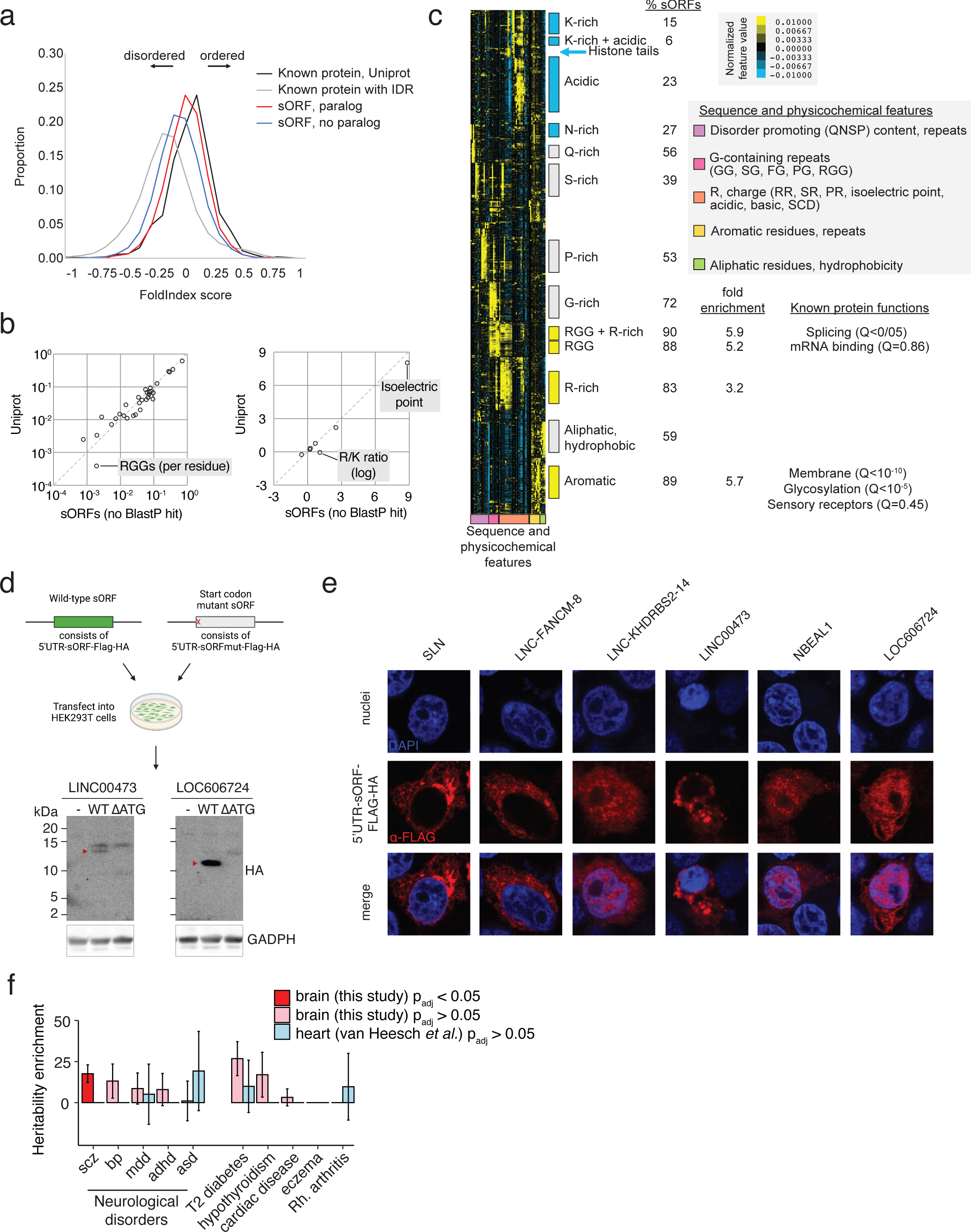
Microprotein functional characterization and disease heritability. (a) FoldIndex score distribution of proteins annotated in Uniprot (black), annotated proteins with intrinsically disordered regions (gray), and sORFs with and without a BlastP hit (red and blue, respectively). (b) Scatterplot of average enrichment per residue of sequence and physicochemical properties in sORFs with no BlastP homology versus annotated proteins (Uniprot). RGG repeats were the most highly enriched of the tested sequence and physicochemical properties in sORFs. (c) Heatmap and hierarchical clustering of z-scores for physicochemical parameters associated with the known disordered proteome (IDRs 21-100 AA in length) as well as sORFs with predicted IDRs that do not have a paralog. Boxes to the right of the heatmap indicate clusters of IDRs with similar properties. Blue = clusters depleted for sORFs, yellow = clusters significantly enriched for sORFs. (d) Western blot of FLAG-HA-tagged unmodified and ATT-mutated *LINC00473* lncRNA, which includes the endogenous 5’UTR. (e) Immunofluorescence of FLAG-HA-tagged *LINC00473* containing the endogenous 5’UTR in HEK293T cells. (f) Heritability enrichment of sORFs in human brain (this study) and human heart (van Heesch 2019) across neurological and non-neurological diseases.

The lack of defined structure may suggest a lack of function for some microproteins; however, many disordered proteins have recently been shown to serve essential cellular functions through a variety of mechanisms, including the tuning of protein interaction specificity and affinity, as well as through the formation of biomolecular condensates ^42–44^. To explore further the potential function of human brain microproteins, we proceeded to compare human brain sORFs with similarly-sized disordered regions from known proteins on the basis of their physicochemical and bulk sequence properties using hierarchical clustering methods ^45^ (Fig. 7c, S7i, Table 4, see Methods). Strikingly, this analysis identified a strong enrichment of brain sORFs (>5x expected, 890 total) in the resulting sequence clusters that were rich in arginine-glycine-glycine (RGG) motifs and/or aromatic residues, as well as, to a lesser extent, clusters rich in arginine residues (>3x expected). Similarly, human brain sORFs were strongly overrepresented (6x expected) in an aromatic amino acid-rich polypeptide cluster. By contrast, clusters of acidic, lysine-rich and polar sequences encompassing known intrinsically disordered proteins were strongly depleted for sORFs, suggesting that brain sORFs do not uniformly cover the entire sequence and function space of known disordered domains but rather display specific sequence features, suggesting likely biological functions. To support this claim, we found that 17 of the human brain microproteins that are predicted to be intrinsically disordered are important for cell survival^9^, further supporting a functional role for these microproteins in the brain.

It is notable that RG and RGG motifs, which can include a single or multiplexed RGG/RG motif ^46, 47^, are known to be important for the regulation of mRNA splicing and translation, and have also been associated with RNA binding and the formation of biomolecular condensates ^46^. Indeed, several previously characterized proteins in RGG- and R-rich clusters are known to interact with RNA in biomolecular condensates (HNRNP/Q, G3BP1/2 and SRSF1) and have been implicated in splicing and mRNA binding by Gene Ontology enrichment analysis (GOrilla, based on the annotations of the proteins that contained intrinsically disordered regions), raising the possibility that the newly identified sORF-encoded microproteins may also interact with RNA-processing complexes to control mRNA splicing, translation, or DNA damage responses in the nucleus. Likewise, the aromatic amino acid-rich polypeptide cluster includes sequences from proteins that are known to be associated with localization in the membrane, glycosylation and functions as neuronal receptors, raising the possibility that some of the newly identified sORFs may encode microproteins with similar functions. Taken together, these analyses thus implicate sORF-encoded brain microproteins in a subset of cellular mechanisms, including mRNA processing and neuronal receptor function.

### In vitro Validation of Candidate Human Brain Microproteins

To independently verify the translation potential and subcellular localization of the identified sORFs, we over-expressed six selected sORFs with their endogenous 5’ UTR and a FLAG-HA epitope tag in heterologous cells (Table 5). These sORFs were all derived from annotated non-coding regions and included both evolutionarily ancient and human-specific sORFs. We confirmed expression of microproteins of expected molecular weight, and subsequent start codon mutation prevented translation (Fig. 7d). These microproteins exhibited a range of subcellular localization patterns (Fig. 7e), suggesting that microproteins expressed in the human brain exhibit diverse cellular interactomes. Expression in heterologous cells thus provides a feasible platform to interrogate the biochemistry of these newly identified human brain microproteins on a candidate basis.

### Human Brain sORFs and Neuropsychiatric Disease

To further investigate the potential of human brain sORFs to regulate critical developmental processes, we tested whether sequence variation associated with brain sORFs might contribute to neuropsychiatric disease risk. We leveraged stratified linkage disequilibrium score regression analysis (see Methods) to determine whether sORF genomic regions contain variants that account for significantly more trait heritability for neuropsychiatric disease than expected by chance. To this end, we assessed heritability enrichment across a number of neurological (schizophrenia, major depression, autism spectrum disorder, bipolar disorder, ADHD, and ALS) and non-neurological (diabetes, cardiovascular disease, rheumatoid arthritis, hypothyroidism, and eczema) conditions for which summary statistics from large GWAS studies are available.

We found that variants within human brain sORFs were significantly enriched for schizophrenia heritability, but not other neurological or non-neurological disorders, after correction for multiple hypothesis testing (Fig. 7f). By comparison, sORFs from the human heart did not demonstrate enrichment for any tested trait (Fig. 7f). Consistent with these findings, the dorsolateral prefrontal cortex, the region used for our Ribo-seq analysis, has been strongly implicated in schizophrenia ^48, 49^, suggesting that sORFs expressed in this region may contribute to schizophrenia disease heritability. While it remains to be determined whether specific sORFs act locally in cis to modulate canonical translation or contribute to this effect via the distinct activities of their encoded microproteins, these findings implicate human variation within these microproteins as a significant contributor to psychiatric disease risk.

## DISCUSSION

RNA translation is a fundamental cellular process that is tightly regulated across human development. The fidelity of translation, stability, and localization of RNA transcripts are critical determinants of brain function, with mRNA translation regulation being a key step that is often mis-regulated in human neurodevelopment and neuropsychiatric disease.^50^ Importantly, studies in other human tissues such as the heart suggest that the translatome is far more complex than previously appreciated ^3–5^, and that the resulting proteome diversity likely contributes to a myriad of functions in these tissues. Remarkably, however, the human brain translatome has remained largely uncharacterized.

We applied ribosome profiling and proteomics to the prenatal and adult human cortex, as well as to hESC-derived neuronal cultures, providing the first large-scale resource of translation events in the developing human brain. By characterizing the translatome across human brain development in >70 individuals, we found that translation is an important mode of regulation for shaping the brain proteome in a way that had not been previously fully appreciated by transcriptome-focused studies. Collectively, our data reveal widespread translation of non-canonical open reading frames in the human brain, including thousands of novel microproteins. As was found to be the case for non-nervous system tissue, we identified in the brain a subset of lncRNAs, uORFs, and other annotated non-coding transcripts that encode translated proteins, some of which we were able to detect by mass spectrometry. In addition to studying the developmental regulation of RNA translation in human brain tissue, we profiled the activity-dependent translatome in hESC-derived neuronal cultures and found that many activity-dependent lncRNAs that were thought to be non-coding are actually translated in this context.

What is unclear for individual ‘non-coding’ transcripts is whether they function in the brain solely as protein-coding RNAs, or whether they have bifunctional potential, whereby the transcript and the encoded protein have independent functions. Recent studies using expression quantitative trait locus analysis suggest that hundreds of lncRNAs have associations with human diseases, and that rare variants in lncRNAs impact complex human traits ^51^. This underscores the importance of discerning the protein-coding potential of brain-expressed lncRNAs, which requires further investigation at the level of individual gene candidates.

The identification of sORFs is a necessary and critical first step in understanding their role in the human brain, and much work remains to understand the function of individual human brain sORFs. To this end, we probed the potential function of these sORFs through analysis of microprotein amino acid sequence, identifying a marked absence of structured domains and likely enrichment for RNA-binding functions. It is tempting to speculate that these disordered microproteins may impact RNA metabolism by enhancing or inhibiting the formation of biomolecular condensates, or by partitioning into them, but this requires extensive experimental validation. Moreover, while the precise mechanisms underlying the emergence of new protein-coding genes are not completely understood, the generation of new ORFs, including sORFs, is likely an important driver of protein evolution. The fact that the majority of sORFs are human-specific renders them interesting candidates in the study of uniquely human features of the brain. The recent evolution, small size, and relatively low translation efficiency of many sORFs also suggests that these sequences represent evolutionary experiments - regions that gain translation capacity in any given lineage but may not always be conserved during further evolution. While these findings may suggest that some newly evolved microproteins are non-functional, we identified many human-specific microproteins that appear to play a key role in cell growth and viability ^9^. In addition, it will be of great interest in the future to understand how sORFs may expand and become fixed in the genome through continued evolution.

It is important to consider several caveats of the current study. First, our ribosome profiling was restricted to bulk tissue measurements, as methods for single-cell ribosome profiling from tissue samples have yet to be reported. In addition, ribosome profiling was largely performed from post-mortem brain tissue, and post-mortem interval-dependent decreases in translation initiation and/or ribosome-RNA binding likely contributed to loss of ORF resolution and thus our experiments may result in an overestimation of truncated ORF annotations. However, overall, the ORFs identified in the present study likely represent an underestimate, as ORFs with substantial ribosomal run-off and loss of periodicity would not be called by the ORF identification algorithms. Notably, we identify many more ORFs in the prenatal brain compared to the adult brain, which is likely at least in part a result of the longer post-mortem interval in adult compared to prenatal samples. Despite these limitations, the finding that the translatome of hESC-derived neurons largely mimics translation in prenatal cortex tissue suggests that our measurements in post-mortem tissue predominantly reflect physiologically relevant translation events. Future studies may leverage cultured hESC-derived neurons to probe the function of sORFs in the context of neuronal function and human disease.

In conclusion, we provide the first large-scale resource for the investigation of translation regulation in the human brain which has yielded unprecedented insight into the under-recognized complexity of the brain translatome and proteome. Importantly, our results identify candidates for future functional characterization of previously unannotated microproteins, opening new opportunities for the investigation of translational regulation in the nervous system and for the elucidation of the function of many new human- and brain-specific microproteins.

## ACKNOWLEDGEMENTS

This research was supported by the Allen Discovery Center program, a Paul G. Allen Frontiers Group advised program of the Paul G. Allen Family Foundation. EED was supported by the Damon Runyon Cancer Research Foundation (DRG-2397-20). ACC was supported by the Hanna H. Gray Fellowship through the Howard Hughes Medical Institute. VL was supported by a Boston Children’s Hospital Career Development Award (to AODL), and VL and NS by a National Human Genomic Research Institute (NHGRI) R01HG010898. AODL and WP were supported by a Manton Center Endowed Scholar Award and the NHGRI U01HG008900. MAS was supported by a National Institute of Mental Health (NIMH) F31MH124393. BTK was supported by a National Institute of Neurological Diseases and Stroke (NINDS) K08 NS112338. MEG was supported by funding from a NINDS R01 NS115965. We thank members of the Greenberg lab for helpful discussions on the manuscript. We thank the Taplin Mass Spectrometry Facility at Harvard Medical School for their technical expertise and analysis of proteomics samples and Dr. Sarah Slavoff for advice on size-selection proteomics. We thank the Broad Institute Genomics Program for next-generation sequencing of ribosome profiling libraries. We are grateful to the NIH NeuroBioBank and the Human Developmental Biology Resource for providing human adult and prenatal brain tissue, respectively. We are grateful to the lab of Dr. Didier Trono for sharing the human transposable element annotation and to Dr. Wade Harper for reagents and technical advice related to iPSC-derived human neurons. We thank the Neurobiology Department and the Neurobiology Imaging Facility for consultation and instrument availability that supported this work. This facility is supported in part by the Neural Imaging Center as part of an NINDS P30 Core Center grant #NS072030. Figures 3d, 4a, and 7d were made with Biorender.

## AUTHOR CONTRIBUTIONS

EED, BTK, and MEG conceptualized the study and designed the experiments. EED and BTK performed ribosome profiling and RNA-seq. EED, BTK, and BF analyzed ribosome profiling and RNA-seq. GC prepared human embryonic stem cell-derived neurons for ribosome profiling. ACC analyzed the insertion of TEs at ORF translation start sites. IP, AA, JF-K and AMM performed physiochemical analysis. VL, AODL, AK, WP, and NS performed phylostratigraphy analysis. BTK prepared samples for proteomics, and BTK, BF, and AK analyzed proteomics data. EED and EGA performed microprotein validation experiments in 293T cells. MS and BB performed disease heritability analysis. EEC and EH provided prenatal brain tissue samples and technical advice for tissue processing. SG assisted with Ribo-seq quality control analyses. EED, BTK, BF, ECG and MEG drafted the manuscript, with input from all co-authors.

## DECLARATION OF INTERESTS

The authors declare no competing interests.

## DATA AVAILABILITY

Human brain primary tissue RNA-seq and Ribo-seq data have been submitted to the database of Genotypes and Phenotypes (dbGaP) under accession number phs002489. NGN2 RNA-seq and Ribo-seq data have been submitted to the Gene Expression Omnibus (GEO) under accession number GSE180240.

---

**Figure S1:**
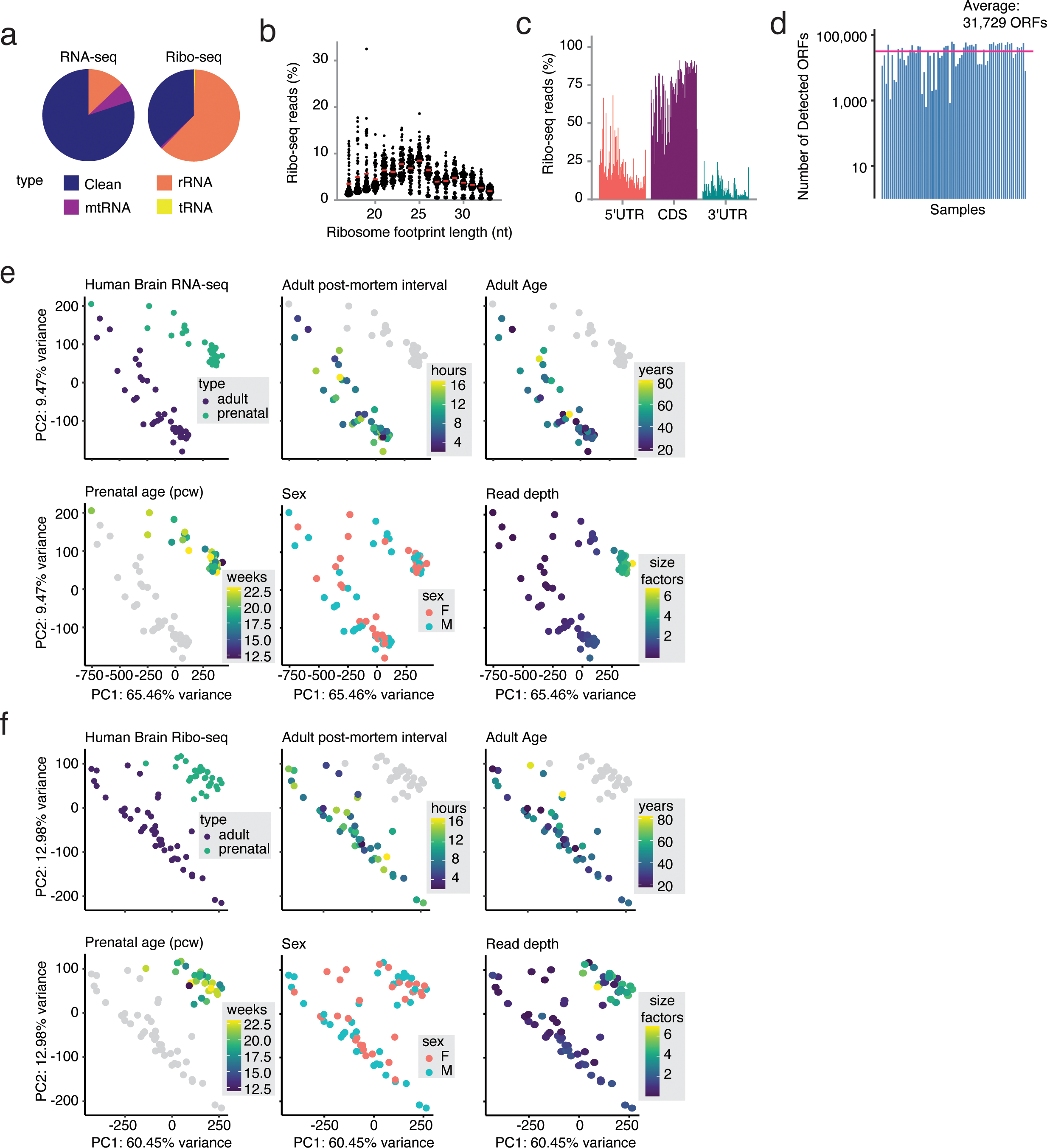
Ribosome profiling captures active translation in the human adult and prenatal brain, Related to Figure 1. (A) Pie chart displaying the fraction of raw sequence reads derived from tRNA, ribosomal RNA (rRNA), mitochondrial RNA (mtRNA), and remaining aligned reads (clean) from human adult and prenatal brain RNA-seq and Ribo-seq. (B) Beeswarm plot of sequenced ribosome footprint lengths across all 73 adult and prenatal brain samples. Red line indicates the average percentage of Ribo-seq reads assigned to a given read length across all samples. (C) Bar plot of the percentage of reads mapping to the coding sequence (CDS) and untranslated regions (5’ and 3’ UTR) of annotated protein-coding genes (Refseq hg38). Each bar represents an individual sample. (D) Bar plot of the number of ORFs identified by RibORF in each sample after filtering (see methods). (E) PCA analysis of all genes in the human brain RNA-seq and (F) Ribo-seq, colored by sample type (adult vs prenatal), post-mortem interval for adult samples, adult age, prenatal age (pcw), sex, and read depth (based on DESeq2 scale factors of estimated library size). The validity of combining samples into two groups in subsequent analyses was confirmed by the finding that these two groups were well separated by PCA analysis for both the transcriptome and translatome.

**Figure S2:**
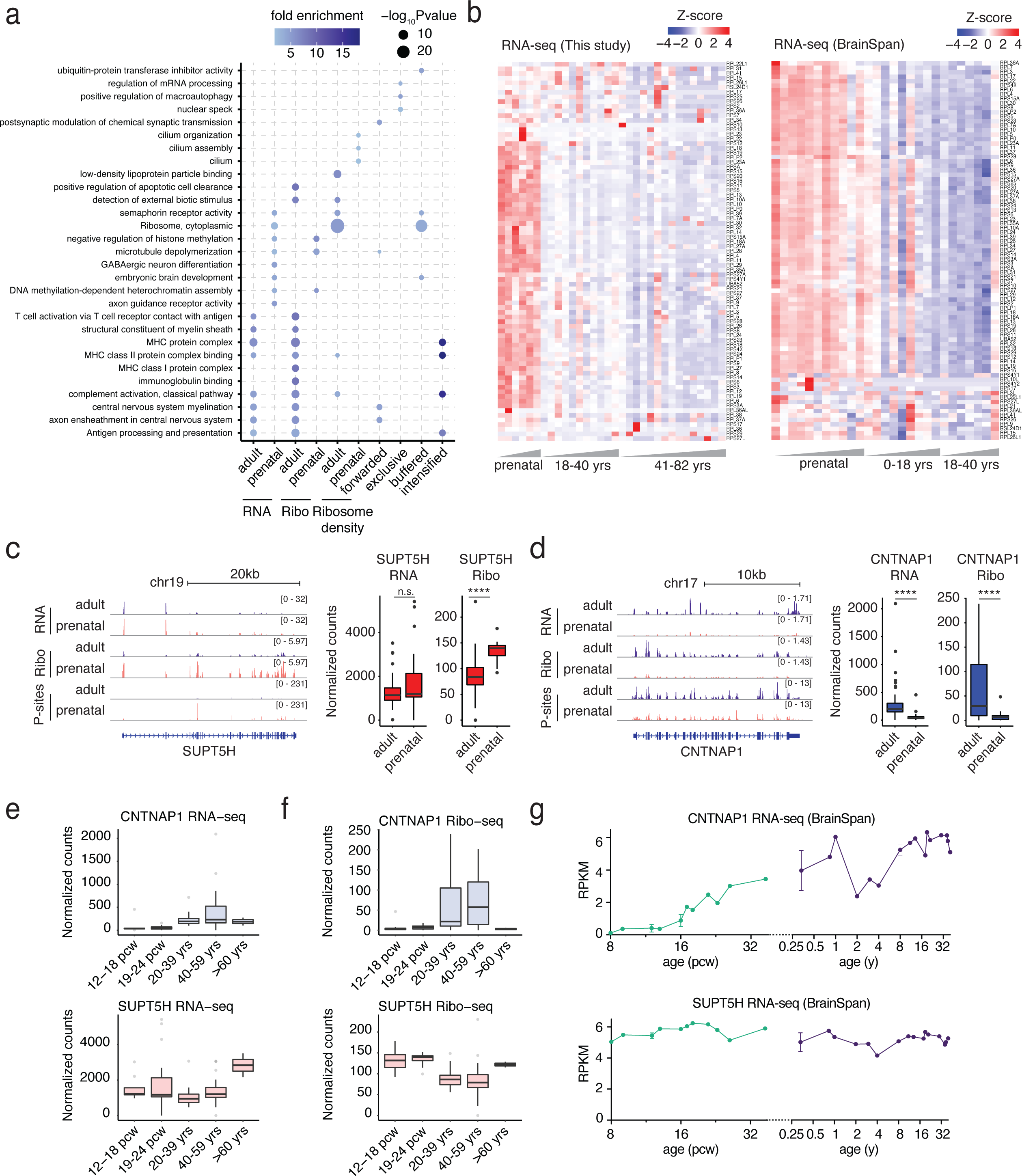
Transcriptional and translational regulation across human brain development, Related to Figure 2. While the translatome has not been previously measured in the developing human brain, our measurements of the transcriptome are consistent with published gene expression data from the BrainSpan Atlas of the Developing Human Brain ^15^ (a) Dot plot of the top enriched GO terms in each regulatory category defined in Fig 2B. (b) Heatmap of RNA-seq expression (row-normalized) for all ribosomal genes in Fig. 2D across all human adult and prenatal samples in this study (left) and in the dorsolateral prefrontal cortex from the BrainSpan Atlas of the Developing Human Brain. (C&D) Genome locus of *SUPT5H*, a transcript regulated only at the level of translation (c), and *CNTNAP1*, a transcript regulated only at the level of transcription (d). (Left) Tracks represent merged and depth-normalized reads across all adult vs. prenatal samples for RNA-seq, Ribo-seq, as well as P-site positions. (Right) DESeq2-normalized reads in adult vs. prenatal samples RNA-seq reads and Ribo-seq P-sites. **** p < 0.05 by DESeq2. (e) Box and whisker plots of DESeq2-normalized RNA-seq reads across human brain samples divided into five age categories (prenatal = 12-18 pcw and 19-24 pcw, adult = 20-39 yrs, 40-59 yrs, >60 yrs) for *CNTNAP1* (left), a translationally forwarded gene, and *SUPT5H* (right), a translationally exclusive gene. (f) Box and whisker plots of DESeq2-normalized Ribo-seq reads across human brain samples divided into five age categories (prenatal = 12-18 pcw and 19-24 pcw, adult = 20-39 yrs, 40-59 yrs, >60 yrs) for *CNTNAP1* (left) and *SUPT5H* (right). (g) Line plot of RPKM values from the BrainSpan Atlas of the Developing Human Brain for *CNTNAP1* (left) and *SUPT5H* (right) across development in the dorsolateral prefrontal cortex.

**Figure S3:**
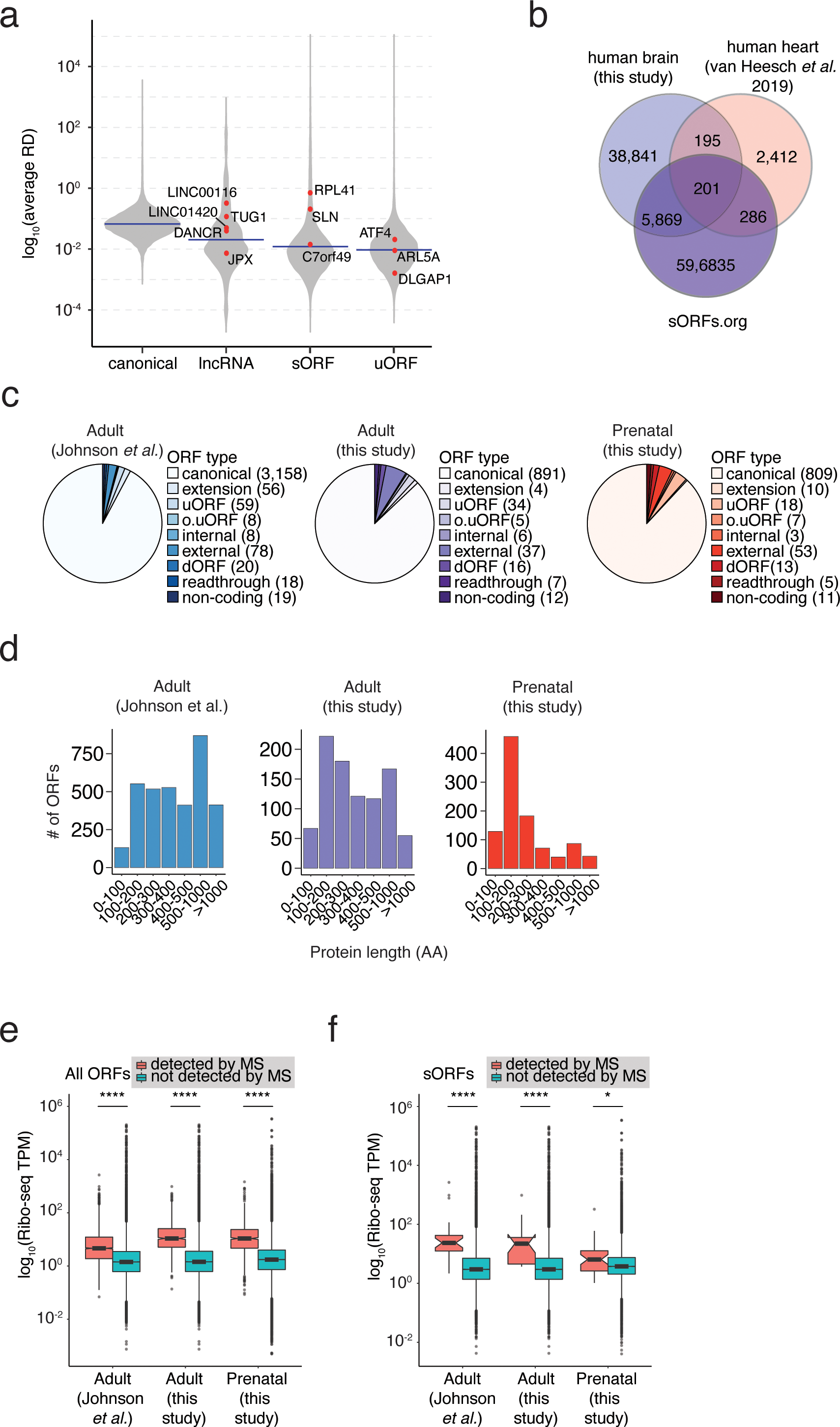
Microprotein expression and validation across brain development, Related to Figure 3. (a) Violin plot of average ribosome density (RD) by ORF type. Previously described ORFs are shown in red. Average ribosome density is shown in blue. (b) Venn diagram of sORFs detected in human brain (this study), human heart (van Heesch *et al*. 2019), or the sORF.org database. In total, 6,468 translated sORFs identified in the human brain perfectly matched the amino acid sequence of a previously reported entry in the sORFs.org database or identified in the human heart, a degree of overlap consistent with prior studies ^4^. (c) Number and type of ORFs identified by size-selection proteomics in the adult and prenatal brain, or by Johnson *et al.* (d) Number of ORFs binned by protein length identified by size-selection proteomics in the adult and prenatal brain, or by Johnson *et al.* (e) Box and whisker plots of Ribo-seq TPM for all ORFs detected by MS and ORFs not detected. (f) Box and whisker plots of Ribo-seq TPM for sORFs detected by MS and ORFs not detected. While only a fraction of the sORFs identified by ribosome profiling were detected by our mass spectrometry analysis, this is not surprising given that such shotgun proteomic approaches have low sensitivity for the detection of individual proteins, particularly if the proteins are transient or low in abundance. Consistent with this finding, the sORF-encoded proteins that we were able to detect by proteomics exhibited a higher average ribosome density compared to all sORFs detected by ribosome profiling.

**Figure S4:**
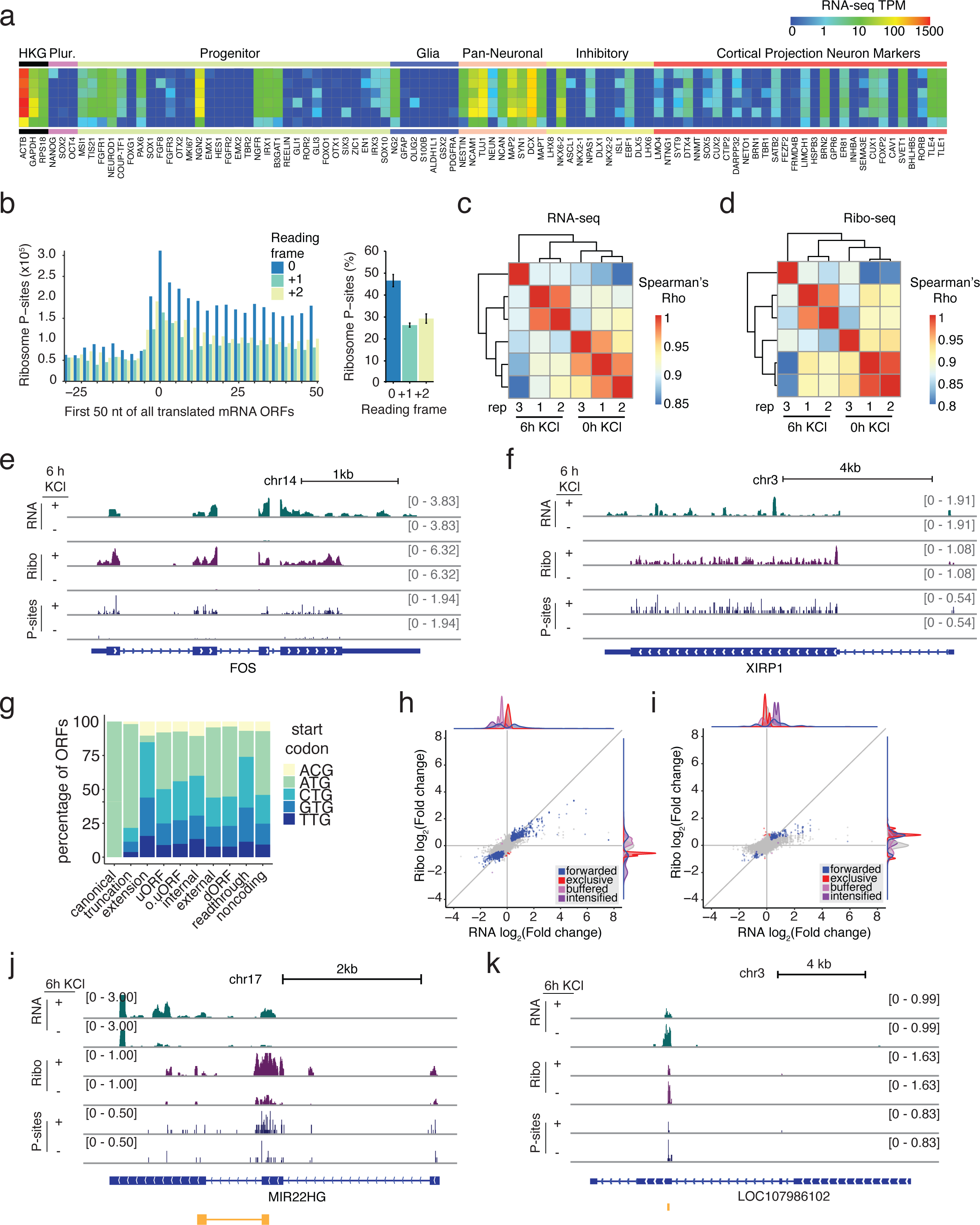
Activity-dependent translation in hESC-derived neurons, Related to Figure 4. (a) Heatmap of RNA-seq TPM from hESC-derived neuronal cultures for marker genes associated with neuronal and non-neuronal cell types. Rows indicate individual samples and biological replicates. The pattern of gene expression observed in these cultures largely mimics the findings in Nehme *et al.* ^28^ (b) Bar plot displaying P-sites derived from offset-corrected Ribo-seq reads in the first 50 nt of annotated ORFs (left) and the aggregate percentage of footprints in each reading frame (right). (c) Heatmap of pairwise Spearman’s r correlation between RNA-seq samples from NGN2-derived neurons for the top 2000 expressed genes. (d) Heatmap of pairwise Spearman’s r correlation between Ribo-seq samples from NGN2-derived neurons for the top 2,000 expressed genes. (E-F) Genomic locus of *FOS*, a classic activity-induced gene in neurons (e), and *XIRP1*, an activity-induced gene that was previously reported in human GABAergic neuron cultures ^52^. (f) Tracks represent merged and depth-normalized reads across 3 biological replicates of membrane-depolarized (6 h KCl) and unstimulated neurons for RNA-seq, Ribo-seq, as well as P-site positions. (g) Breakdown of translated ORFs of each type identified in NGN2-derived neurons. (h&i) We found that activity-dependent changes in ORF translation were largely driven by transcriptional changes rather than a shift in ribosome density for both canonical ORFs and sORFs. This finding is consistent with observations that activity-dependent translation events coupled to transcription are transient and likely return to basal levels within six hours of membrane depolarization. (h) Scatterplot of fold-changes between stimulated and unstimulated neurons for all canonical ORFs in Ribo-seq data and the corresponding gene in RNA-seq data. Translationally forwarded genes (blue), exclusive genes (red), buffered genes (light purple), and intensified genes (dark purple) are highlighted. (i) Scatterplot of fold-changes between stimulated and unstimulated neurons for all sORFs in Ribo-seq data and the corresponding gene in RNA-seq data. Translationally forwarded genes (blue), exclusive genes (red), buffered genes (light purple), and intensified genes (dark purple) are highlighted. (j&k) Genomic loci of two activity-dependent lncRNAs with evidence of translation, *MIR22HG* (j) and *LOC107986102* (k). Tracks represent merged and depth-normalized reads across 3 biological replicates of membrane-depolarized (6h KCl) and unstimulated neurons for RNA-seq, Ribo-seq, as well as P-site positions. sORF identified by RibORF is shown in gold.

**Figure S5:**
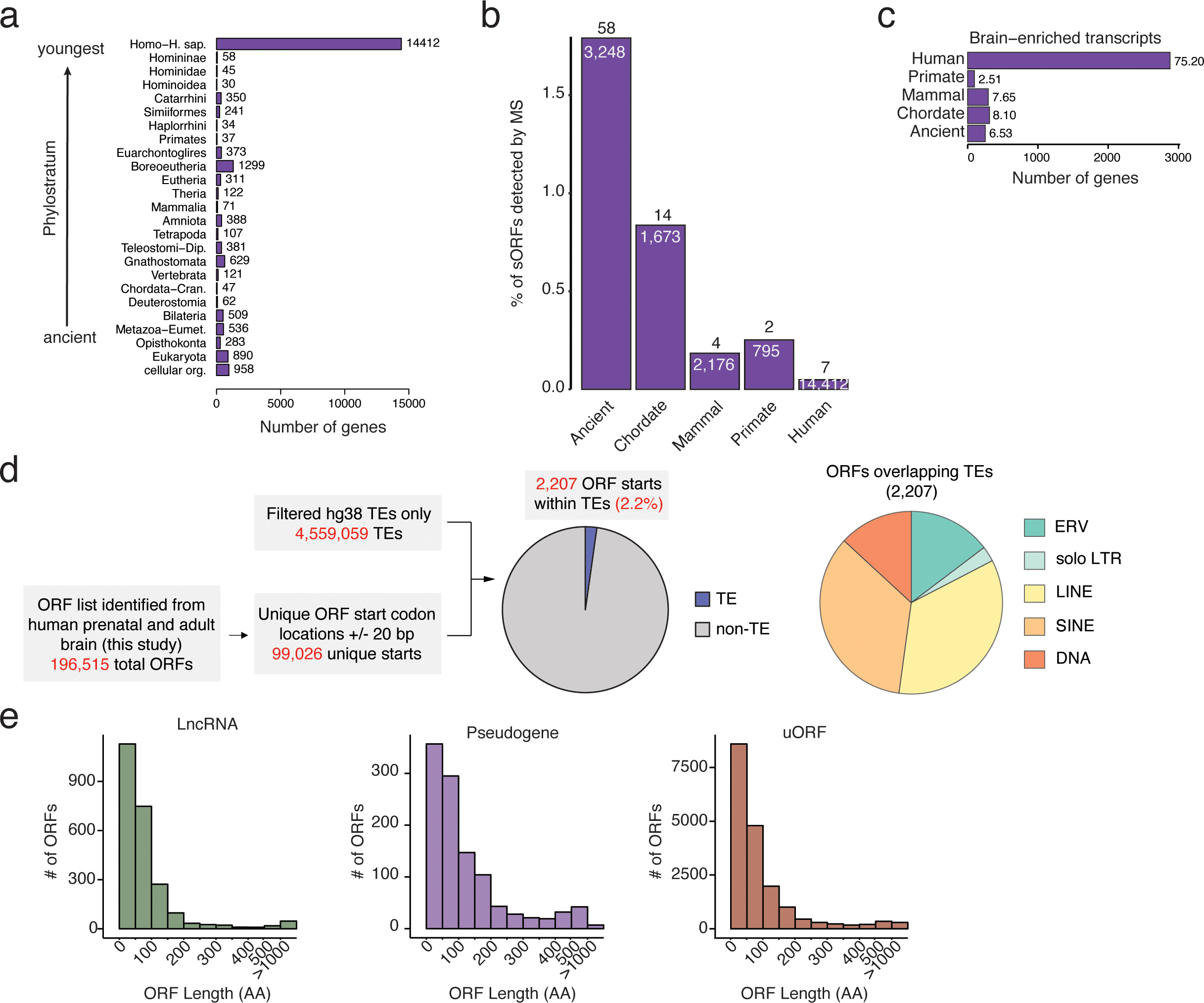
Evolutionary origins of human brain sORFs, Related to Figure 5. (a) Bar plot of the number of sORFs (40-110 AA) grouped by evolutionary age. (b) Bar plot of the number of sORFs (40-100 AA) detected by mass spectrometry (See Fig. 4a) grouped by evolutionary age. (c) Number and percentage of sORFs ≥40 AA that are translated from brain-enriched transcripts, grouped by evolutionary age. (d) Criteria for filtering TE insertion events at start codons (left) and pie chart of TE type for all ORFs in our dataset with a TE insertion at the start codon. (e) Histogram of ORF length for all ORFs encoded within lncRNAs, pseudogenes, and uORFs.

**Figure S6:**
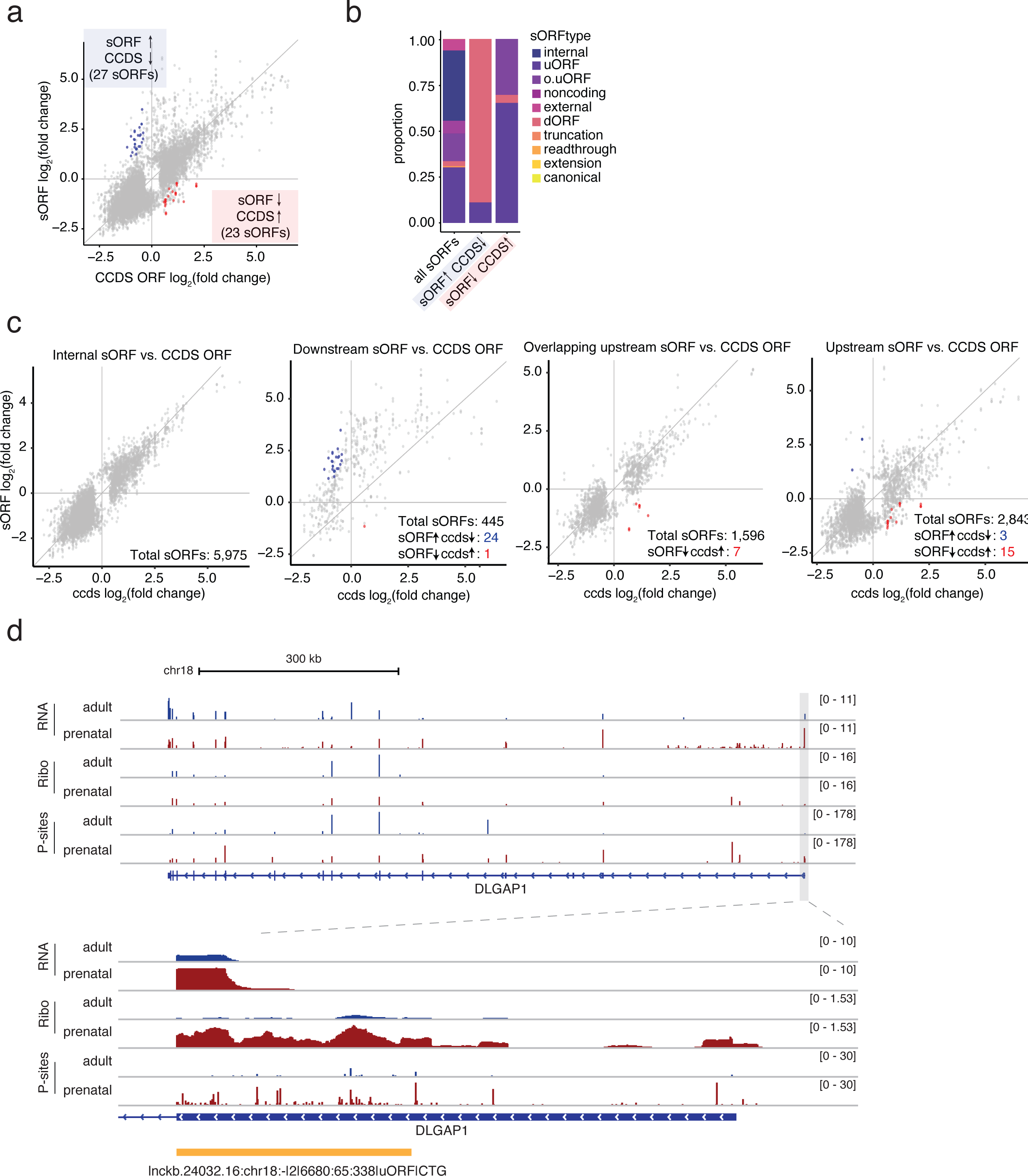
Effects of uORF expression on downstream ORF translation, Related to Figure 6. (a) Scatterplot of fold-changes in translation between adult and prenatal brain for sORFs and canonical ORFs expressed from the same transcript. Positive values indicate enrichment in the adult brain, whereas negative values indicate enrichment in the prenatal brain. Red points indicate genes where sORF translation is significantly (p_adj_ < 0.05) enriched in prenatal brain while canonical ORF translation is significantly (p_adj_ < 0.05) enriched in adult brain. Blue points indicate genes where sORF translation is significantly (p_adj_ < 0.05) enriched in adult brain while canonical ORF translation is significantly (p_adj_ < 0.05) enriched in prenatal brain. **While most sORFs exhibited concordant translation with their associated canonical ORFs across development, we identified 50 sORFs that were discordant with nearby canonical ORF translation, and these discordant sORFs were strongly enriched for uORFs translated from 5’UTRs of annotated protein-coding genes. (b)** Stacked bar plot of numbers and percentages of sORFs detected in human brain (all sORFs), or sORFs exhibiting oppositely regulated expression across development compared to a canonical ORF translated from the same gene, separated by sORF type. **(c)** Scatterplot of fold-changes in translation between adult and prenatal brain for sORFs and canonical ORFs expressed from the same transcript, separated by type of ORF. Positive values indicate enrichment in the adult brain, whereas negative values indicate enrichment in the prenatal brain. Red points indicate genes where sORF translation is significantly (p_adj_ < 0.05) enriched in prenatal brain whereas canonical ORF translation is significantly (p_adj_ < 0.05) enriched in adult brain. Blue points indicate genes where sORF translation is significantly (p_adj_ < 0.05) enriched in adult brain whereas canonical ORF translation is significantly (p_adj_ < 0.05) enriched in prenatal brain. (d) Genomic locus of *DLGAP1*. Tracks represent merged and depth-normalized reads across all adult vs. prenatal samples for RNA-seq, Ribo-seq, as well as P-site positions. The sORF identified by RibORF is shown in gold.

**Figure S7:**
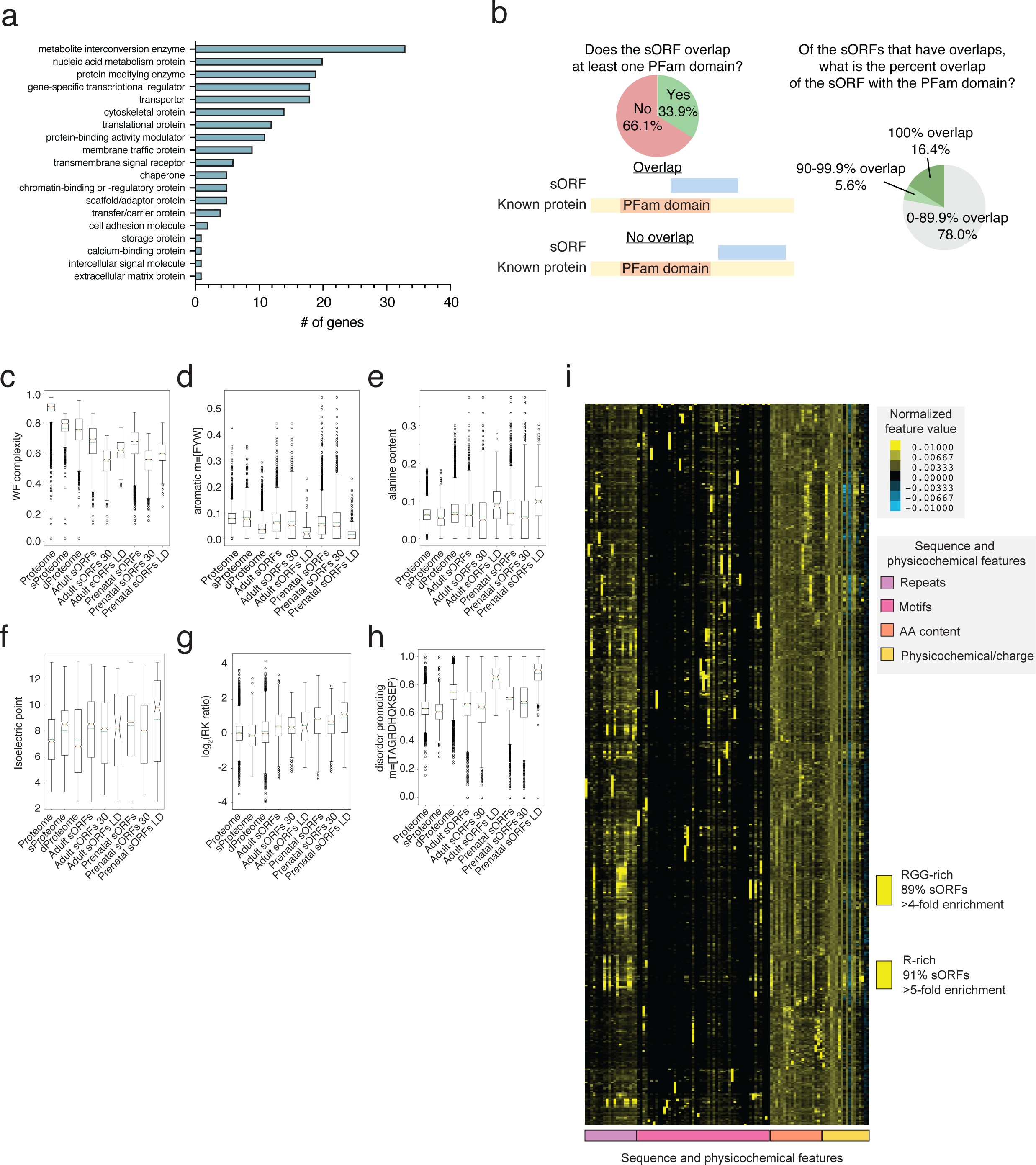
Microprotein functional characterization and disease heritability, Related to Figure 7. (a) Protein functions of known genes that share homology with sORFs. (b) Characterization of sORFs that overlap with a domain in the PFam database. (c) Box and whisker plot of sequence complexity in the known human proteome (Proteome), sORFs (sProteome), the disordered proteome (dProteome), adult and prenatal sORFs (adult sORFs and prenatal sORFs), adult and prenatal sORFs smaller than 30 AAs (adult sORFs 30 and prenatal sORFs 30) and adult and prenatal sORFs with long intrinsically disordered regions (adult sORF LD and prenatal sORF LD). (d) Box and whisker plot of the proportion of aromatic amino acids (phenylalanine, tyrosine, and tryptophan) in the categories of ORFs described in c. (e) Box and whisker plot of the proportion of alanine in the categories of ORFs described in c. (f) Box and whisker plot of the isoelectric point for the categories of ORFs described in c. (g) Box and whisker plot of the arginine to lysine ratio in the categories of ORFs described in c. (h) Box and whisker plot of the proportion of disorder-promoting amino acids (threonine, alanine, glycine, arginine, asparagine, histidine, glutamine, lysine, serine, glutamate, and proline) in the categories of ORFs described in c. (i) Heatmap and hierarchical clustering of z-scores for 109 sequence and physicochemical features associated with the known disordered proteome as well as all sORFs that do not contain a BlastP hit. Boxes to the right of the heatmap indicate clusters of IDRs with similar properties. Yellow = clusters significantly enriched for sORFs.

## METHODS

### Human brain samples

All human tissue research was approved by the Harvard Medical School Institutional Review Board. Adult brain tissue samples were obtained from the National Institute of Health (NIH) NeuroBioBank. Prenatal brain samples were obtained from the Human Developmental Biology Resource (HDBR) and the University of California San Francisco Pediatric Neuropathology Laboratory.

### Ribosome Profiling of Human Brain Tissue

Ribosome profiling was performed using a protocol modified from McGlincy et al.^53^ Frozen brain tissue (∼80 mg) was thawed on ice. Each sample was dounced 15x in 400 µL ice cold lysis buffer: 20 mM Tris pH 7.4, 150 mM NaCl, 5 mM MgCl2, 1 mM DTT, 100 µg/mL cycloheximide (Sigma). The lysate was further sheared using a 26-gauge syringe. The lysate was clarified by centrifugation at 20,000 x g for 10 minutes at 4°C. Supernatant (10 µL) was removed, added to 300 µL Trizol, and frozen at -80°C for future RNA-seq library preparation. RNA concentration in the remaining supernatant was quantified using RNA Qubit. Lysate (30 µg) was subjected to RNase I digestion (0.5 U RNaseI per µg RNA) at room temperature for 45 minutes with gentle agitation.

After RNase digestion, 10 µL SuperasIN (Thermo) was added to each sample, and the samples were transferred to ice. To isolate ribosome protected fragments, the RNase-digested lysate was transferred to Ultra-clear 11x34 mm centrifuge tubes (Beckman Coulter) and underlaid with 0.9 mL sucrose cushion. Samples were centrifuged in a TLS-55 rotor at 51,000 rpm for 2 hours at 4°C. The supernatant was discarded and the pellet was resuspended in 300 µL TRIzol. Ribosome-protected fragments were purified from TRIzol using the Zymo Direct-zol kit. RNA was precipitated by adding 38.5 µL RNase-free water, 1.5 µL Glycoblue, 10 µL 3 M sodium acetate pH 5.5, and 150 µL isopropanol to 100 µL eluted RNA. The mixture was incubated overnight at -20°C. Samples were centrifuged for 30 minutes at 20,000 x g at 4°C. The supernatant was discarded, and the RNA pellet was resuspended in 5 µL 10 mM Tris pH 8. Five µL 2X denaturing sample loading buffer (980 µL formamide, 20 µL 500 mM EDTA, 300 µg bromophenol blue) was added to each sample, then the sample was denatured at 80°C for 90 seconds. Ribosome-protected fragments, along with control oligos^53^, were run on a 15% polyacrylamide gel at 200 V with 12 µL NEB miRNA marker. The gel was stained with SYBR gold in 1X TBE. Gel fragments between 17 and 34 nucleotides were excised and placed in a microfuge tube with 400 µL gel extraction buffer. Samples were frozen on dry ice for 30 minutes, then thawed overnight with gentle agitation.

After overnight gel extraction, 400 µL eluate was transferred to a new microfuge tube. The RNA was precipitated by adding 1.5 µL glycoblue and 500 µL isopropanol. After overnight incubation at -20°C, the sample was centrifuged at 20,000 x g for 30 minutes at 4°C. The supernatant was discarded, and precipitated RNA was resuspended in 4 µL 10mM Tris pH 8. Samples were then dephosphorylated using T4 PNK (4 µL RNA in 10 mM Tris pH 8, 0.5 µL T4 PNK enzyme, 0.5 µL T4 PNK buffer, and 0.5 µL Superasin) at 37°C for 1 hour. Samples were then subjected to SPRI clean up: 50 µL of sample in RNase-free water was added to 90 µL RNAclean beads and 270 µL isopropanol. After washing with 85% ethanol, beads were resuspended in 7 µL RNase-free water. The supernatant was collected, and we proceeded with next-generation sequencing library preparation using the Clontech smRNA library prep kit according to the manufacturer’s instructions. Libraries were sequenced on an Illumina NovaSeq S2 with single-end 1x50 nt reads. Samples were always processed in large batches of a maximum of 24 samples to avoid sample processing biases.

### Human Neuron Differentiation

The use of hESCs was approved by the Harvard Medical School Embryonic Stem Cell Research Oversight (ESCRO) Committee. We used the H9 NGN2 hESC line, harboring a doxycycline-inducible NGN2 construct in the *AAVS1* locus, referred to here as H9 NGN2^54^. We collected human neurons from three independent differentiation cohorts, and each replicate exhibited characteristic gene expression patterns reported previously (Fig. S6A).^28^ On the day prior to cell harvest, neurons were silenced with TTX and APV, which antagonize sodium channels and NMDA receptors, respectively. H9 NGN2 cells were cultured in mTeSR Plus media (STEMCELL Technologies) on tissue culture plates coated with hESC-qualified matrigel (Corning). They were passaged using Dispase (1 mg/mL, Life Technologies) until ready for differentiation. A previously published protocol that combines developmental patterning and NGN2 induction was adapted to differentiate H9 NGN2 into neurons.^28^ At day 0, cells were treated with Accutase (StemPro Accutase, Life Technologies) and plated in single cells at 50,000 cells/cm^2^ in mTeSR Plus media supplemented with 10 µM Y-27632 (STEMCELL Technologies) on tissue culture plates coated with 336.67 µg/mL Growth Factor Reduced matrigel (Corning). On day 1, the medium was replaced with KSR media (Knockout DMEM medium, 15% knockout serum replacement (KOSR), 2 mM L-Glutamine, 1X MEM non-essential amino acids (MEM NEAA), 1X penicillin/streptomycin (pen/strep) and 1X 2-mercaptoethanol (all Gibco)) supplemented with 100 nM LDN193189, 2 µM XAV939 (STEMCELL Technologies), 10 µM SB431542 hydrate and 2 µg/mL doxycycline hyclate (Sigma). Day 2 media was 50% KSR media/50% NIM media supplemented with LDN/XAV/SB and 2 µg/mL doxycycline. NIM media consisted of DMEM/F-12 medium, 1X GlutaMAX, 1X MEM NEAA, 1X pen/strep, 0.16% D-glucose (Sigma) and 1X N2 supplement-B (STEMCELL Technologies). Day 3 media was NIM media supplemented with 2 µg/mL doxycycline. At day 4, cells were treated with Accutase and plated in single cells at 40,000 cells/cm^2^ in NB media (Neurobasal medium (without glutamine), 1X GlutaMAX, 1X MEM NEAA, 1X pen/strep and 1X N2 supplement-B) supplemented with 1X B27 without Vitamin A, 2-2.4 µg/mL mouse laminin (Gibco), 1 µM ascorbic acid, 2 µM dibutyryl cyclic-AMP (Sigma), 20 ng/mL brain-derived neurotrophic factor, 10 ng/mL glial-derived neurotrophic factor (rhBDNF and rhGDNF, Peprotech), 10 µM Y-27632 and 2 µg/mL doxycycline on the tissue culture plates coated with 336.67 µg/mL Growth Factor Reduced matrigel. The media at day 4 without Y-27632 and doxycycline is referred to as complete NB (cNB) media. On day 5 the media was replaced with cNB media. Thereafter, half of the media was replaced weekly with cNB2x, where concentrations of all the supplements to the NB media (except Y-27632) were doubled. In between each media change, media was directly supplemented with 2 µg/mL doxycycline on the third day of the week (days 8, 15 and 22). Cells were silenced on day 27 with TTX and APV, which antagonize sodium channels and NMDA receptors, respectively. Cells were collected at day 28 in 1X PBS supplemented with 1X cycloheximide after being stimulated with 55 mM KCl for 0 or 6 hours.

### Ribosome Profiling of Human Neurons

Ribosome profiling of human NGN2-induced neurons was performed as described above for human brain tissue, except that RNase I digestion time was 15 minutes.

### RNA-seq library preparation

RNA-seq libraries were prepared from 10 ng total RNA using the SMARTer Stranded Total RNA-seq Pico Input Mammalian V2 kit (Takara Bio) according to the manufacturer’s instructions. Samples were multiplexed with Illumina TruSeq HT barcodes and sequenced on a NextSeq 2000 with single-end 1x75 nt reads. Samples were always processed in large batches of a maximum of 24 samples to avoid sample processing biases.

### Analysis of RNA sequencing data

In an effort to capture the most complete picture of translation, including the potential translation of brain-specific lncRNAs, RNA-seq and Ribo-seq reads were mapped to the lncRNA knowledge base (lncRNAKB) annotation.^55^ This annotation includes experimental evidence of lncRNA expression across 31 solid human normal tissues, including the brain, providing a comprehensive resource of transcripts and transcript isoforms in the human brain. Sequencing reads were aligned using Hisat2 (version 2.1.0) to the *H. sapiens* genome (GRCh30) and transcriptome (lncRNAKB). Alignments and analysis were performed on the Orchestra2 high performance computing cluster through Harvard Medical School. Aligned bam files were sorted using Picard Tools (version 2.8.0), stranded bedGraphs were generated using STAR, and reads were quantified over annotated exons using HTSeq-count (version 0.9.1).

### ORF calling and filtering with RibORF

Sequencing adapters were removed using Cutadapt (version 1.14), trimmed fastq files were aligned to hg38 ribosomal RNA sequences, and unaligned reads were mapped to the hg38 genome and lncRNAKB transcriptome^55^ using STAR (version 2.7.3a) with standard settings and the following modified parameters: *--clip5pNbases 3, --seedSearchStartLmax 15, -- outSJfilterOverhangMin 30 8 8 8, --outFilterScoreMin 0, -outFilterScoreMinOverLread 0.66, - outFilterMatchNmin 0, -outFilterMatchNminOverLread 0.66, --outSAMtype BAM Unsorted*. Aligned bam files were sorted using Picard Tools (version 2.8.0) and stranded bedGraphs were generated using STAR. The RibORF pipeline was run on each sample individually using standard parameters. Due to template switching during library preparation, reads contained three untemplated bases at the 3’ end that were not included in the alignment but added to the length of each read. Therefore, reads 30-33 nt in length (corresponding to RNA fragments 27-30 nt) were analyzed for three-nucleotide periodicity within known protein-coding ORFs (RefSeq). For each sample we selected only the read lengths for which at least 50% of the reads matched the primary ORF of known protein-coding genes in a meta-gene analysis. Samples with fewer than two read lengths passing filtering were removed from further analysis. Read lengths were offset-corrected and RibORF was used to predict ORFs with a minimum length of 8 AA and translation probability >0.7.

After running the RibORF pipeline on each brain sample individually, information from RibORF output files was used to generate GTF and BED files for all ORFs identified in each sample. ORFs with lengths of zero and ORFs annotated as non-coding despite being detected in protein coding genes were eliminated. Using Bedtools version 2.27.1 and the GRCh38 primary assembly human genome file, DNA sequences were associated with each exon of each remaining ORF. ORFs that did not end in stop codon sequences (“TGA”,”TAA”,”TAG”) were eliminated. Using the R library micropan version 2.1, DNA sequences for each complete ORF were translated into protein sequences. Of note, ORFs with start codons “GTG” or “TTG” are translated with a Methionine as the initial amino acid despite these sequences not typically encoding methionine in other positions in a protein-coding DNA sequence, per existing literature on non-canonical start codon usage in translation. Finally, all remaining ORF information was collapsed into one table, and duplicate ORFs, defined as ORFs in the same genomic position with identical protein sequences, were eliminated. When eliminating duplicates, ORFs identified of the most common ORF type identified by RibORF were conserved, according to the following order of priority, from highest to lowest: canonical, truncation, extension, overlap, uORF, internal, external, polycistronic, readthrough, and non-coding ORFs. ORFs annotated as type “seqerror” were eliminated. After combining ORF outputs from all samples, ORFs that were only detected in one sample were eliminated. After the removal of singleton ORFs, duplicate ORFs were once again eliminated according to the same priority scheme, leaving only one entry for each ORF that was detected in at least two samples in the dataset. The LncRNAKB annotation was used to assign a specific ORF type to each ORF. In order to be designated a sORF, an ORF had to be 100 amino acids or less in length, and not fully overlap in-frame with a canonical protein-coding ORF.

### Differential expression and GO enrichment analysis

RNA-seq gene expression was quantified as described above, and lowly expressed genes were filtered for counts per million > 1 in at least 2 samples using edgeR (version 3.26.8). Ribo-seq expression was quantified by counting the number of P-sites over a given ORF. To identify differences in transcription and translation between adult and prenatal human brain, two-way differential expression analysis was performed using deltaTE^17^ in R 4.0.1. Read normalization and size factor estimation were performed on RNA-seq and Ribo-seq data simultaneously, and ORF types were subsetted for display purposes. GO enrichment analysis was performed using gProfiler2 in R (version 0.2.0), with a custom background of expressed genes based on expression-filtered RNA-seq genes and FDR < 0.05.

### uORF/canonical ORF correlation analysis

For each mRNA transcript detected in each individual human brain sample, upstream open reading frame sequences identified by RibORF were joined to produce one singular sample-specific upstream open reading frame region. For each of the upstream open reading frame regions, raw counts were generated by quantifying total P-sites across each region, from which TPMs were calculated. These TPMs were compared to canonical open reading frame translational efficiency values to characterize the relationship between upstream open reading frame utilization and canonical open reading frame translational dynamics at the level of individual genes.

### Protein Sequence Analysis by LC-MS/MS

Size-selected proteomics of the human adult and prenatal brain, as well as hESC-derived neurons, was performed at the Taplin Biological Mass Spectrometry Facility at Harvard Medical School. Excised gel bands were cut into approximately 1 mm^3^ pieces. Gel pieces were then subjected to a modified in-gel trypsin digestion procedure. Gel pieces were washed and dehydrated with acetonitrile for 10 min, followed by removal of acetonitrile. Pieces were then completely dried in a speed-vac. Rehydration of the gel pieces was with 50 mM ammonium bicarbonate solution containing 12.5 ng/µL modified sequencing-grade trypsin (Promega, Madison, WI) at 4°C. After 45 min, the excess trypsin solution was removed and replaced with 50 mM ammonium bicarbonate solution to just cover the gel pieces. Samples were then placed in a 37°C room overnight. Peptides were later extracted by removing the ammonium bicarbonate solution, followed by one wash with a solution containing 50% acetonitrile and 1% formic acid. The extracts were then dried in a speed-vac (∼1 hr). The samples were stored at 4°C until analysis.

On the day of analysis, samples were reconstituted in 5 - 10 µL of HPLC solvent A (2.5% acetonitrile, 0.1% formic acid). A nano-scale reverse-phase HPLC capillary column was created by packing 2.6 µm C18 spherical silica beads into a fused silica capillary (100 µm inner diameter x ∼30 cm length) with a flame-drawn tip. After equilibrating the column, each sample was loaded via a Famos auto sampler (LC Packings, San Francisco CA) onto the column. A gradient was formed, and peptides were eluted with increasing concentrations of solvent B (97.5% acetonitrile, 0.1% formic acid).

As peptides eluted, they were subjected to electrospray ionization and then entered into an LTQ Orbitrap Velos Pro ion-trap mass spectrometer (Thermo Fisher Scientific, Waltham, MA).

Peptides were detected, isolated, and fragmented to produce a tandem mass spectrum of specific fragment ions for each peptide.

### Mass Spectrometry Analysis

Thermo-Fisher raw files were loaded into MaxQuant version 1.6.17.0 for the peptide search. Each file corresponded to one brain sample and was labeled as its own experiment in the search. Default parameters, including specific trypsin digestion, methionine oxidation and protein N-terminal acetyl variable modifications, and carbamidomethyl-fixed modifications were used. We uploaded a custom protein FASTA file for our search using the protein sequence identified in our RibORF post-processing. For adult brain mass spectrometry, we used a protin FASTA file containing only sequences from adult samples that passed our quality control metrics, and the same for prenatal brain mass spectrometry. In each case, “truncation” type ORFs were excluded because of their redundancy to canonical protein sequences. The protein search in MaxQuant was run using an Amazon Web Services client to optimize speed and efficiency. Only peptides with a score >50 were considered for subsequent analysis.

### Physiochemical analysis

sORFs were searched using Blastp (Version 2.6.0, -evalue 0.0001, -word_size 4) against a database of all protein translations from Gencode v29 (https://www.gencodegenes.org/human/release_29.html; downloaded on Aug 2, 2019). Locations of significant hits were then compared to PFam annotations (from Ensembl accessed through Ensembl API) and any sORF with at least one residue overlap was considered overlapping. Although we allowed any degree of overlap, we found that many of the sORFs had near complete overlap with PFam domains (Fig. S4D). FoldIndex score^41^ is defined as 2.785*hydropathy - abs(net charge) - 1.151. Physicochemical and sequence properties of sORFs and IDRs were computed using custom python codes (https://github.com/IPritisanac/mol.feat.idrs.git). All analyses of sORFs, IDRs and reference proteins were performed on protein sequences between 21 and 100 amino acids. 27,110 sORFs, 19,652 IDRs and 908 Uniprot human reference proteins met this criterion. Normalization, filtering and clustering of sequence properties was performed using cluster3.0^56^ with the following parameters: median centering of columns, normalization of columns, retaining sequences with at least 3 observations with absolute value>0.01 and weighting columns using default options and clustering using average linkage hierachical clustering. This process left 16,905 sequences in the cluster analysis of which 6,910 were IDRs and 10,095 (59%) were sORFs. Clusters were visualized and selected manually using treeview v1.1.6r4^57^ and enrichment analysis was performed by selecting the IDRs from each cluster and using the 6910 IDRs in the entire cluster analysis as the background set. These lists were entered into the GOrilla webserver.^58^ The enrichment or depletion of sORFs in each cluster was computed by comparing the ratio of sORFs in each cluster to the expected ratio of 10,095/6,910.

### Phylostratigraphy analysis

All ORFs with an amino acid length of ≥40 amino acids were analyzed as described previously^5^, using TimeTree^59^ to identify the minimal evolutionary age for every protein-coding gene. The evaluation is based on sequence similarity scored with Blastp and identifying the most distant sequence in which a sufficiently similar sequence appears. Each protein sequence was used to query the non-redundant (nr) NCBI database with a Blastp e-value threshold of 10e-3 and a maximum number of 200,000 hits. We identified the phylostratum in which each ORF appeared. Each phylostratum corresponds to an evolutionary node in the lineage of the species, as listed in the NCBI Taxonomy database. For clarity, we aggregated results into the following evolutionary eras: Ancient (phylostrata 1-7, from cellular organisms through Deuterostomia (290 – 747 millions of years ago (Mya))), Chordates (phylostrata 8-17, from Chordata through Amniota (747 - 320 Mya)), Mammals (phylostrata 18 - 22, from Mammalia through Euarchontoglires (320 - 91 Mya)), Primates (phylostrata 23-29, from Primates through Homininae (91 – 6.6 Mya)), and Humans (phylostrata 30-31, including Homo sapiens, 6.6 Mya to present).

### Transposable element insertion at start codons

To identify ORFs whose start codons derive from transposable elements, we intersected our ORF start codons with a TE annotation kindly provided by the lab of Dr. Didier Trono.^60^ First, we created a list of all start codons in different categories (protein-coding, lncRNA, uORF, sORF, pseudogene) by collapsing all ORFs that share a start position. We extended this start codon position to a 10 bp window and intersected this with a bed file of all TEs in the human genome using BedTools. For any ORF start codon that overlapped a TE, we used a table of TE subfamily ages from Dfam to estimate the oldest possible lineage in which that TE may exist in the human genome.^32, 61^

### Microprotein overexpression and western blot

To test the translatability of sORFs, dsDNA sequences containing the sORF endogenous pseudo 5’UTR (defined as the upstream DNA sequence from the sORF start codon), sORF protein sequence, and a FLAG-HA tag in-frame with the sORF protein sequence was synthesized by Genscript and cloned into an FUGW overexpression vector (Addgene #14883). A negative control construct in which the start codon was mutated to an ATT was generated using a Quikchange II Site-directed mutagenesis kit (Agilent 200521). The wild-type and mutant plasmids were verified by Sanger sequencing. The designed sequences used in this study are listed in Table 5.

Both wild-type and mutant plasmids were transfected into HEK293T cells using Lipofectamine 3000 reagent (Invitrogen). 24h later, cells were harvested and resuspended in RIPA buffer (Sigma) supplemented with protease inhibitor cocktail (Roche). Protein concentration was measured by Bradford assay, and 20 µg protein lysate was denatured at 95°C for 5 min and then separated on a 10-20% Tris-tricine gel (Invitrogen) at 125V for 90 min. Proteins were transferred to a nitrocellulose membrane (Bio-Rad) at 115V for 90 min, and the membrane was blocked with 5% non-fat dry milk in TBST for 1h. Membranes were incubated with anti-HA primary antibody (1:1000) (CST) in 5% non-fat dry milk in TBST overnight at 4°C. Membranes were washed 4x in TBST at room temperature then incubated with secondary antibodies conjugated to IRdye 800 (1:10,000) and imaged with LiCOR Odyssey.

### Immunofluorescence

HEK293T cells were grown on glass slides for 24h and transfected as described above. Cells were fixed with 4% paraformaldehyde for 30 min at room temperature and washed three times with ice-cold PBS. The cells were permabilized and blocked for 1h at room temperature using 5% donkey serum in PBST (1X PBS, 0.1% Triton X-100). Coverslips were incubated with anti-FLAG mouse primary antibody (1:1000) (Sigma-Aldrich) overnight at 4°C. Coverslips were washed 3x in PBST at room temperature then incubated with fluorescently-labeled secondary antibody (1:2000, Alexa Fluor 488 anti-mouse) for 1h at room temperature. Coverslips were washed 3x in PBST at room temperature and mounted onto superfrost glass slides using DAPI Fluoromount-G (Thermo Fisher Scientific). Images were visualized using a LEICA SP8 confocal microscope using a 63x objective.

### Disease heritability enrichment

Heritability enrichment analysis was performed using stratified linkage disequilibrium score regression (LDSC)^62^ as described in Boulting et al..^52^ Briefly, we obtained a baseline model of 54 annotations from Finucane et al. and augmented the model with three human brain-specific regulatory annotations from Rizzardi et al..^63^ We then tested heritability enrichment of neurological and non-neurological diseases for annotations defined from sORFs from 3 categories: all sORFs in the human brain or NGN2-derived neurons, all sORFs in the human heart^4^, and any sORF translated from an activity-dependent RNA. When comparing enrichment of an annotation across traits, we corrected for multiple hypothesis tests using the Holm step-down procedure. The studies from which summary statistics were obtained for each tested trait are provided in Table 6.

## Notes

### Competing Interest Statement

The authors have declared no competing interest.

